# Computational Properties of the Visual Microcircuit

**DOI:** 10.1101/2020.07.30.229435

**Authors:** Gerald Hahn, Arvind Kumar, Helmut Schmidt, Thomas R. Knösche, Gustavo Deco

**Author notes:** Corresponding author: Gerald Hahn, Center for Brain and Cognition, Computational Neuroscience Group, Department of Information and Communication Technologies, Universitat Pompeu Fabra, C/Roc Boronat 138, 08018 Barcelona, Spain.

## Abstract

The neocortex is organized around layered microcircuits consisting of a variety of excitatory and inhibitory neuronal types which perform rate-and oscillation based computations. Using modeling, we show that both superficial and deep layers of the primary mouse visual cortex implement two ultrasensitive and bistable switches built on mutual inhibitory connectivity motives between SST, PV and VIP cells. The switches toggle pyramidal neurons between high and low firing rate states that are synchronized across layers through translaminar connectivity. Moreover, inhibited and disinhibited states are characterized by low- and high frequency oscillations, respectively, with layer-specific differences in frequency and power which show asymmetric changes during state transitions. These findings are consistent with a number of experimental observations and embed firing rate together with oscillatory changes within a switch interpretation of the microcircuit.

## Introduction

The neocortex is a recurrent network of morphologically diverse inhibitory interneurons and excitatory pyramidal neurons (PYR)^1–4^. The majority of interneurons can be assigned to biochemically defined classes, such as parvalbumin (PV), somatostatin (SST), and vasoactive intestinal polypeptide (VIP) positive cells^5^. These neurons are distributed across layers and connected according to an intricate circuit diagram with intra- and interlaminar connections^6–10^. The discovery of regularities within the connectivity pattern of excitatory and inhibitory neurons prompted researchers to propose the existence of canonical microcircuits^11,12^, which implement elementary computations that are repeated across the brain^13^.

To identify such computations, research has focused on a better description of the functional role of individual neuron types by selective optogenetic activation and silencing of specific cell types^5,14–16^. These studies have not only highlighted an essential role of inhibitory neurons to balance excitation, but also recognized disinhibitory subcircuits which release pyramidal neurons from strong inhibition^15,17–21^. Moreover, neuronal oscillations in different frequency bands have been attributed to the activity of different interneuron types^22–24^. While the function of simplified circuits with multiple interneuron types, attributed to the superficial cortical layer, have been investigated theoretically ^25–29^, the dynamics of more complex networks comprising multiple layers with translaminar connectivity remain unexplored. Moreover, it is unclear how firing rate descriptions of microcircuit function relate to oscillatory behavior of cortical networks, which can differ across cortical layers ^30–32^. This is crucial to interpret meso- and macroscopic signals from LFP, EEG or MEG recordings in light of circuit function, where access to firing rate information is not possible.

To address these questions, we take a computational approach and isolate the role of different neurons in rate- and oscillation based functioning of the layered microcircuit of the primary mouse visual cortex, for which the most comprehensive connectivity diagram to date is available^7^. Modeling permitted to go beyond of what is possible with available experimental tools and we not only characterized the effect of selective activation/suppression of different neuron types, but also perturbed specific connections and test their impact on microcircuit dynamics and response properties.

We found that the superficial and deep layers in the visual microcircuit can operate in two different states, each with different excitation-inhibition balance: Inhibition dominated state controlled by SST neurons and a disinhibited state governed by PV neurons. By perturbing connections of different types of interneuron we confirmed that disparities in recurrent connections within these inhibitory cell classes play a crucial role for the different EI-balance in the two states. Two mutual inhibitory motifs that include SST, PV and VIP cells serve as ultrasensitive or bistable switches with different sensitivity, which can toggle the microcircuit between the two states. Such a state change in one layer can propagate through translaminar connections to the other layer. Notably, we also found that in the inhibited regime beta-band oscillations were more prevalent especially in the deep layer, whereas in the disinhibited state gamma oscillations emerged predominately in the superficial layer, similar to experimental observations^30,32^ We also provide a mechanistic explanation to other empirical findings such as asymmetric changes in oscillation power and frequency during state transitions as seen with the presentation of visual stimuli with increasing size^24,33^. Thus, our results provide a comprehensive description of state-dependent effects of different inhibitory interneuron types with testable predictions and relate rate and oscillation –based accounts of microcircuit functioning.

## Results

In this study, we investigate the computational properties of a detailed microcircuit of the mouse visual cortex^34^ (Fig. 1a). This network consists of two different layers (superficial and deep), representing L2/3 and L5 of the primary visual cortex and each containing four different cell types that are connected within and between layers: excitatory pyramidal cells (PYR), and three different classes of inhibitory cells (PV, SST and VIP). The connectivity was corrected for the different prevalence of each cell type^34^ and scaled with a global parameter (*G*) to approximate effective coupling changes. The variable *G* could be related to the overall cell count in the microcircuit^35,36^ (Fig.1b). The population dynamics of each neuron type was given by a firing rate model (see Methods). This microcircuit model also displayed noisy oscillations, which allowed us to study both firing rate and oscillatory behavior as measured by variations in power and frequency of the local field potential (LFP), approximated by the rate of the pyramidal cell populations. We first examined spontaneous interactions across all neurons and then drove specific neuron classes, simulating input from remote cortical and subcortical sources.

**Figure 1.**
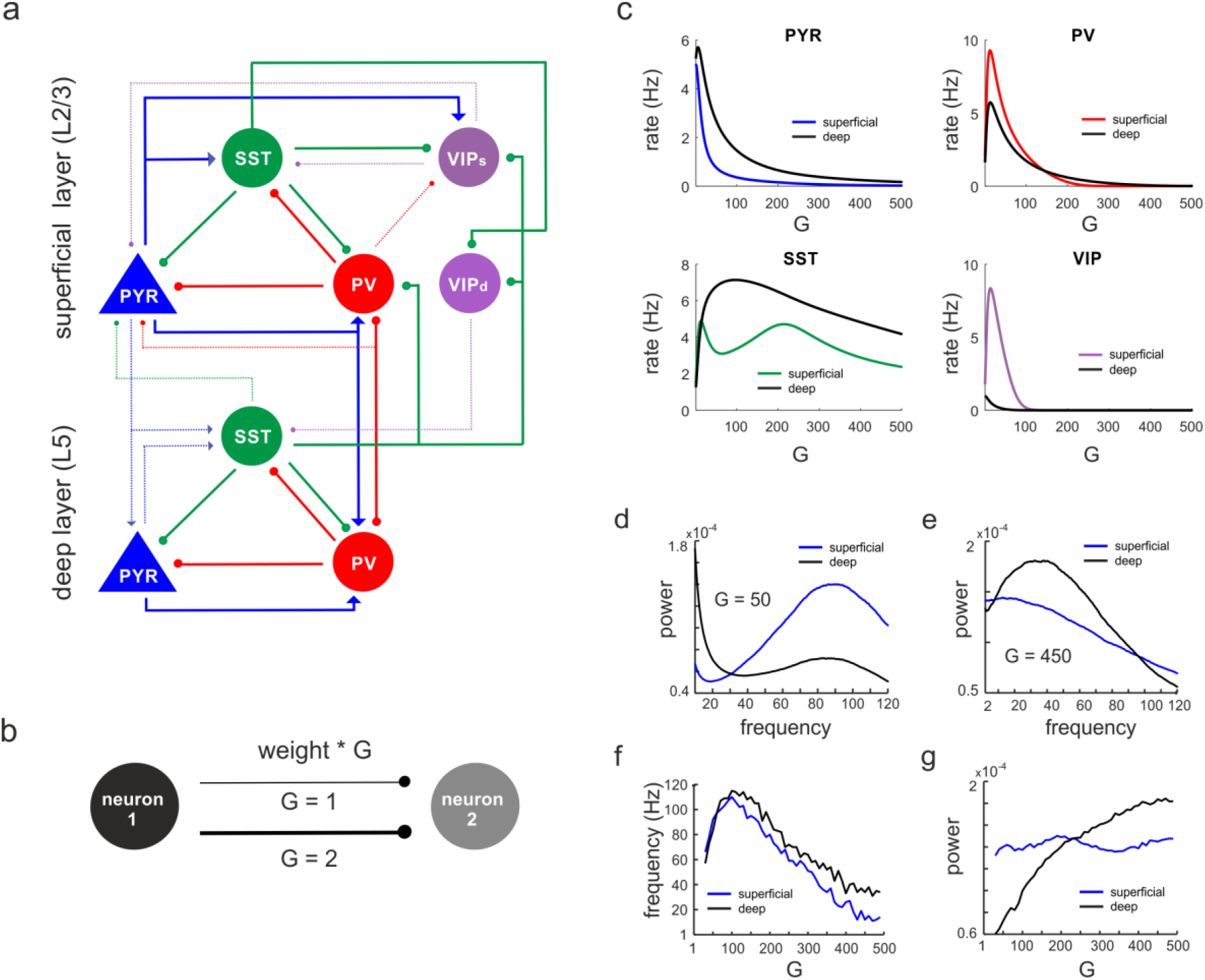
Network anatomy and spontaneous activity. **a)** Layout of the local network with a superficial layer that includes 4 different cell types in layer 2/3 of the mouse visual cortex and three cell types in a deep layer representing L5. Even though residing in the superficial layer, the VIPd cell type was functionally associated with the deep layer, as it mainly innervates L5. The connectivity strength (w) is represented by the thickness of the lines. Solid lines: w > 0.1, dashed lines: intermediate weights: 0.04 > w < 0.1, weak weights (w < 0.04) are not shown. **b)** Schematic showing the scaling of a connection by a coupling parameter *G*. **c)** Mean spontaneous rate for all cell types in superficial and deep layers as a function of the coupling parameter *G* **d-e)** Example power spectra of LFP in superficial and deep layers for two different values of *G*. **f-g)** Frequency and power of oscillatory peaks in LFP spectra as a function of *G* for both layers.

### Spontaneous activity

First, we systematically scaled the microcircuit connectivity by *G*, and measured the steady-state firing rates of all neurons without any external input. A sharp increase in pyramidal, PV and VIP cell activity with *G* was followed by a rapid decrease of mean rates in both layers (Fig. 1c). SST neuron also behaved similar to PV and VIP cells but both rise and decay in their activity was much slower. The average firing rates of pyramidal and SST neurons was higher in the deeper layer, in accordance with experimental results in mice^37,38^. Power spectral analysis of the LFP showed a clear peak, whose frequency and power varied with the coupling parameter in a layer specific manner (Figs. 1d-e). Generally, frequencies first increased within the high gamma range from ~60 Hz (for small *G*) to ~110 Hz (for *G* = 100) across both layers (Fig. 1f). For *G* > 100, the dominant LFP frequency steadily decreased to a low gamma (~40 Hz, superficial layer) or low beta range (~15Hz, deep layer) for *G* = 500. Interestingly, the frequency consistently remained higher in deep layers, consistent with recent experimental findings in mouse V1^31^. Moreover, high gamma frequencies (G < 250) were stronger in superficial layers, whereas the power of slower oscillations (G > 250 was higher in deep layers (Fig. 1g), again in congruence with experimental studies ^32,39–41^.

### Origin of different firing rate and oscillation in deep and superficial layers

Next, we investigated the anatomical origin of firing rate and oscillation differences across layers by modifying specific connections. We targeted three connections, which show pronounced asymmetry across layers: PYR^sup^→PYR^deep^ connection, translaminar projections of SST cells, and recurrent inhibition among PV neurons (PV-PV connections) (Fig. 2a, Supplementary Table 4). The removal of the PYR^sup^→PYR^deep^ connection strongly reduced the firing rate differences across the layers (Fig. 2b). Removal of the translaminar SST connection which only projects from deep to superficial layer had a smaller impact on the firing rate difference. Moreover, the disinhibitory PV-PV connections are considerably stronger in deep layers (Supplementary Table 4). When we changed the PV-PV connections such that their strength was the same in the deep and superficial layers, the firing rate difference between the two layers was also reduced. Modifying all three connections simultaneously almost completely abolished rate inequality between layers. Likewise, differences in oscillation power in different frequency bands across layers were suppressed (Figs. 2c-d, compare with Figs. 1d-e). Thus, our model suggests that stronger excitation and disinhibition in the deep layer together with more inhibition in the superficial later underlie the experimentally observed firing rate and oscillation power differences between deep and superficial layers.

**Figure 2.**
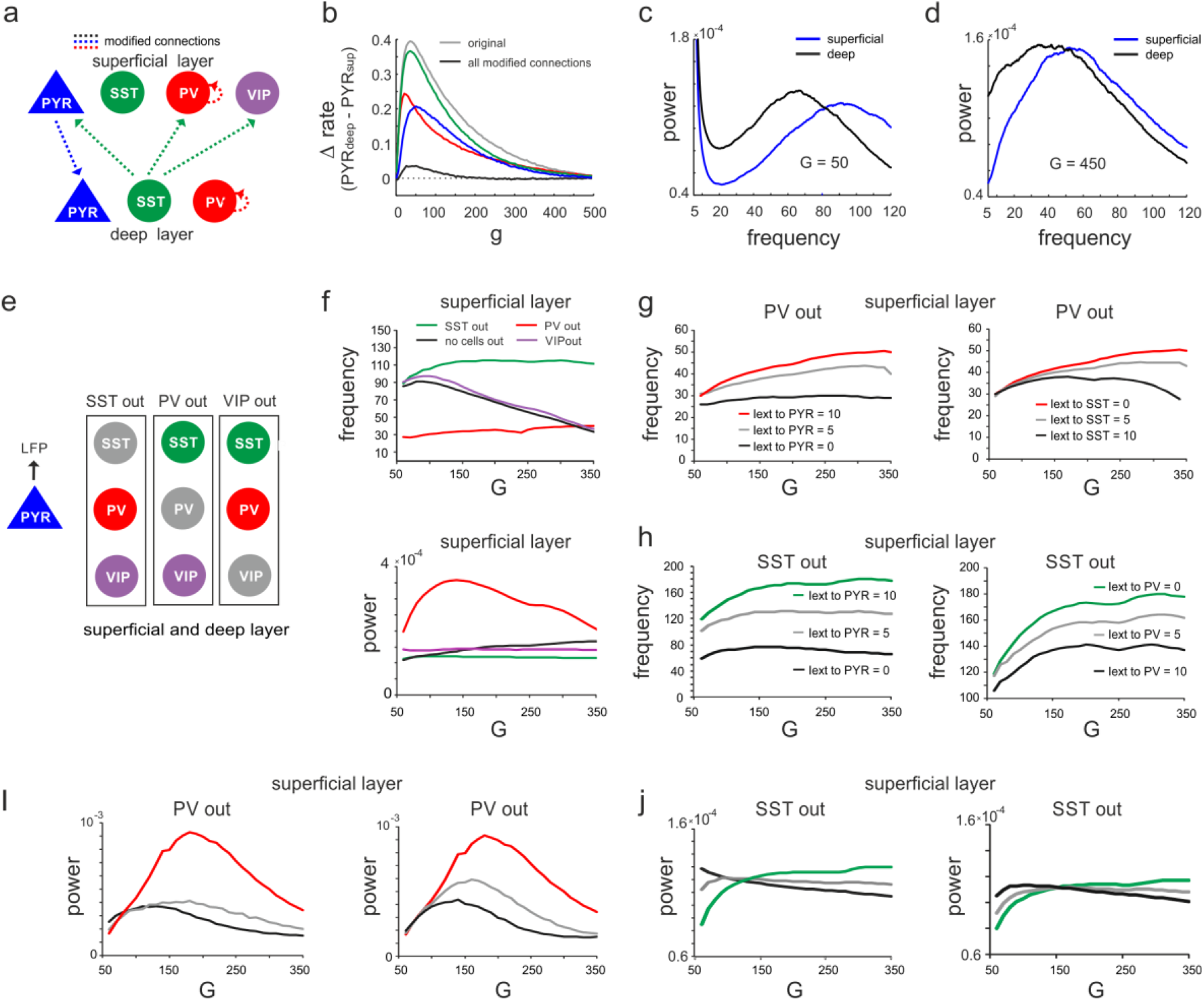
Effect of connectivity lesions on rate and spectral properties across layers. **a)** Schematic two layer network. The blue and green translaminar connections were removed, while the weights of the red connections were both set to the same value of the deep recurrent PV connection. **b)** Effect of connectivity modification on rate difference between superficial and deep layer as a function of *G*. **c-d)** Power spectra of both layers for two values of *G* after all connectivity modifications were applied. **e)** Diagram depicting three different cell lesion simulations. Gray circles: connections of this cell type to all other cells were set to zero. **f)** Peak LFP frequency (top) and power (bottom) as a function of *G* for different cell lesions in the superficial layer. **g)** Peak frequency in the superficial layer as a function of *G* after PV cell inactivation and different levels of input to PYR (left) or SST cells (right). **h)** Superficial layer peak frequency after SST lesion as a function of *G* and varying input to PYR or PV cells. **i-j)** Same as in **g-h)** for oscillatory peak power in the superficial layer.

### Effect of silencing specific inhibitory cells

The above results suggest how different inhibitory neurons may cause changes in the dominant oscillation frequency and power. Notably, relative dominance of SST cells should be accompanied by low frequency oscillations owing to their slow synaptic time constants. By contrast, relative dominance of PV neurons should give rise to high frequency oscillations given their faster time constant. To test this hypothesis, we silenced individual inhibitory cell types in both layers by removing all their connections and examined the effect on oscillation frequency and power of the LFP (Fig. 2e). Knocking out PV cells was accompanied by a slow oscillation (~30 Hz) in both layers with a frequency that remained approximately stable with *G*. By contrast, when SST cells were removed the network oscillated at high frequency (~110 Hz) across all tested values of *G* (Fig. 2f, top, Supplementary Fig. 2a, left). Note that this increase in the oscillation frequency was not due to disinhibition from SST silencing, because PYR firing rate decreased in both knock-out cases with *G* (Supplementary Fig. 1), while the frequency remained high. In accordance with the hypothesis, for small values of *G* when PV rate is high, the oscillation frequency in the intact network approached the frequency seen in the SST knock-out case. As *G* was increased and SST activity surpassed PV firing rates, the oscillation frequency decreased and converged to values seen in the PV knock-out scenario. Importantly, oscillatory power was overall higher in the PV knock-out case with low frequency and lower in the SST silenced network with faster oscillations across all tested values of *G* (Fig. 2f, bottom, Supplementary Fig. 2a, right). In contrast to PV and SSP cells, VIP cell silencing only slightly increased the oscillation frequency, but did not influence the relative decrease in frequency as a function of *G*. However, when we manipulated frequency and power in the knock-out networks, we found that they changed symmetrically. In both PV and SST silenced circuits, driving PYR cells or the remaining inhibitory cell jointly increased or decreased power and frequency in superficial and deep layer, except for the deep layer after SST silencing (Figs. 2g-j, Supplementary Fig. 2b-c). Thus, the relative dominance of PV and SST cell activity is an important factor that determines the oscillation frequency and power, which are inversely related in the full model, but positively correlated in the partly silenced network.

### Two different states and state switching dynamics of the microcircuit

Thus far, we changed the relative prevalence of given interneuron types by scaling the connectivity matrix or silencing individual cell types or connections. Visual inspection of the microcircuit revealed two prominent mutual inhibition motifs. SST cells exhibit reciprocal inhibitory connections with PV cells and also VIP cells in each layer, which brings these cell pairs in competition with each other (Fig. 1a). We hypothesized that driving one inhibitory cell type will functionally silence competing cells and toggle the circuit between different inhibitory states, which may differ in terms of PYR firing rate and their oscillation profile. To verify this hypothesis, we first tested whether input to VIP or PV cells can suppress SST activity. To this end we enhanced the SST activity by injecting additional input to SST cells in both layers (I_ext_ = 5Hz). Next, we stimulated either VIP or PV cells in both layers simultaneously, mimicking feedforward input from layer 4 to PV cells or feedback input from upstream areas to VIP cells, which may target superficial and deep layers simultaneously^32,42^. Responses were measured from PYR and SST cells for different values of *G* (Figs. 1a-b, see Supplementary Figs. 3a-b for responses of other cell types).

When VIP cells were driven (Supplementary Fig. 3a), we found two main effects. First, VIP connections to SST cells suppressed SST rates, whereas PYR and PV cells increased their firing across both layers, because they were released from SST inhibition, in particular for higher *G* values (Fig.3a and Supplementary Fig. 3a). However, as VIP input increased, PYR and PV cells were gradually suppressed again in the superficial layer, while their firing rate was only slightly affected in the deep layer. This was caused by the inhibitory connection from VIP to PYR cells in the superficial layer and indeed its removal resulted in a response similar to the deep layer two layers (Fig. 1a, Supplementary Fig. 4).

**Figure 3.**
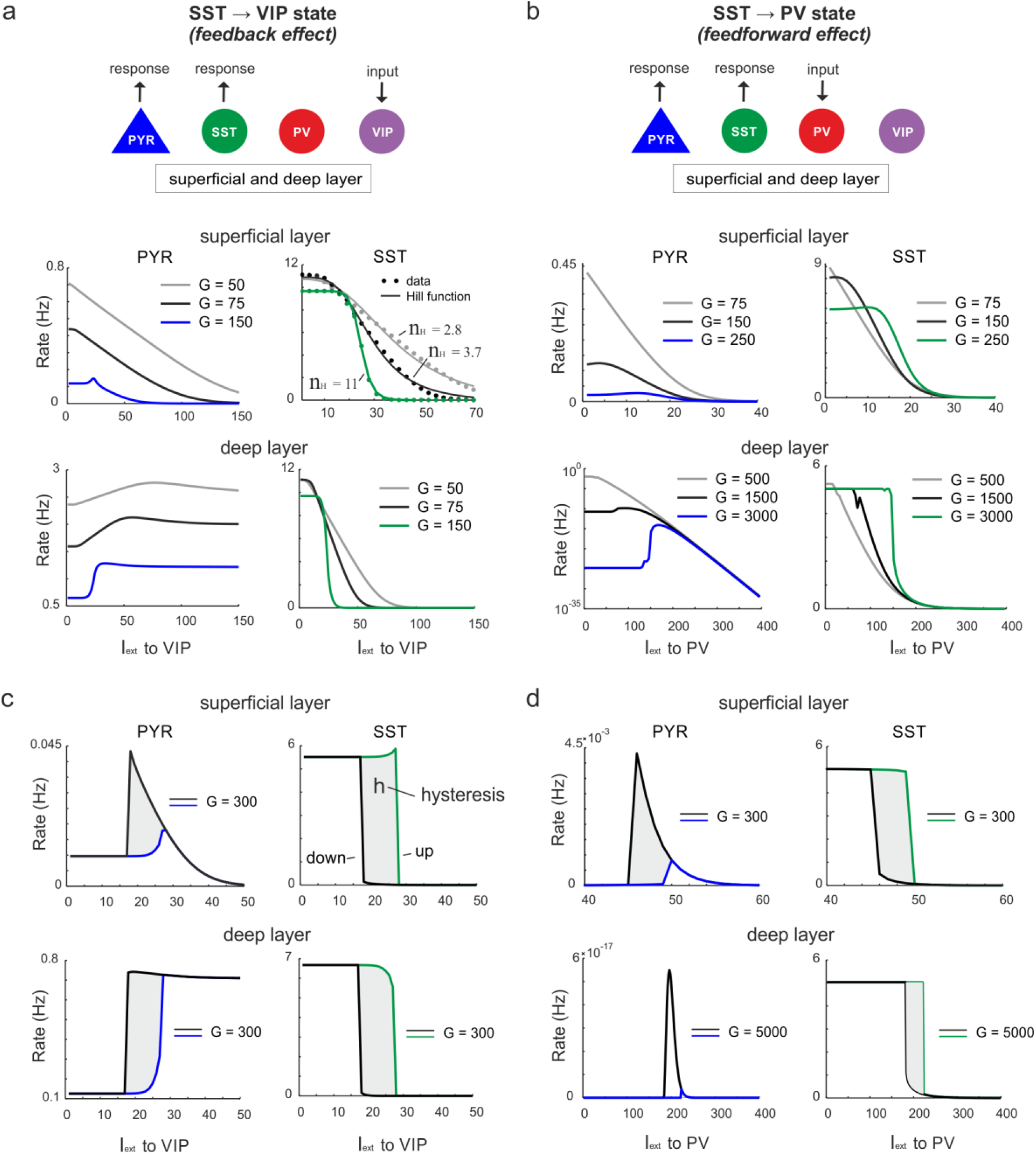
Ultrasensitivity and hysteresis in the visual microcircuit. **a)** Response of PYR and SST cells in superficial and deep layers after input to superficial and deep VIP cells (top), mimicking cortical feedback. SST cells were driven with a constant input I_ext_ = 5 to enhance SST activity. Responses are shown for three different values of *G* (bottom). SST response curves were fitted with the Hill function, yielding a different Hill coefficient (nH) for each curve. **b)** Same as in a) for simultaneous input to superficial and deep PV cells, simulating feedforward inhibition. **c)** PYR and SST cell response to increasing (up branch, in blue/green) and decreasing input (down branch, in black) to VIP cells for an exemplary value of g and both layers. Note that g is higher than in a). The network displays hysteresis in each layer (shaded region h). **d)** Same as in c) for simultaneous input to PV cells in both layers.

Second, as *G* increased, the responses of SST cells became more switch like with sigmoidal curves in both layers, a hallmark of an ultrasensitive switch in many biological systems^43^ (Fig. 3a). A similar sigmoidal decrease in SST firing rates with initial PYR disinhibition followed by inhibition was observed in both layers when only PV cells were driven (Fig. 3b, Supplementary Fig. 3b). However, when PV cells were stimulated, switch like ultrasensitive responses occurred at larger *G*-values as compared to VIP input, especially in the deep layer. By contrast, VIP cell induced inhibition on PYR cells was weaker than PYR suppression mediated by PV cells (Fig. 3a-b, Supplementary Fig. 3c).

In some biological systems ultrasensitivity with sigmoidal response curves is accompanied by bistable behavior^44^, characterized by state transitions that are not reversed when the input is withdrawn. A telltale sign of bistability is the presence of hysteresis, that is the response curves change as a function of the direction in which the state change was triggered. To test for bistability in the microcircuit, we first applied an increasing current to VIP or PV cells in both layers simultaneously, followed by current in the decreasing direction for different values of the coupling parameter. We found that hysteresis appeared at sufficiently high *G*-values, as visible by the appearance of non-congruent response curves for the up and down direction in all cell types of both layers for VIP (Fig. 3c, Supplementary Fig. 5a) and PV input (Fig. 3d, Supplementary Fig. 5b). Hysteresis in the deep layer for PV input required very strong input, even though a small hysteresis effect was observed for smaller *G* that was transmitted from superficial layers via translaminar connections (Supplementary Fig. 5c).

Inhibition-based ultrasensitivity and hysteresis commonly require strong inhibitory interactions between the components of the system^44^. Therefore, we hypothesized that the enhanced mutual inhibitory connections between SST < > VIP and SST< >/PV cells due to the scaling by *G* underlie the sigmoidal and hysteretic response curves. Indeed, selectively increasing these weights was sufficient for ultrasensitivity and hysteresis to appear in the microcircuit (Supplementary Fig. 6).

Next, to quantify the switching behavior of the microcircuit we fitted Hill function to the SST firing rate response curve (see Methods) and estimated the Hill coefficient (n_H_, Fig. 3a), a measure for (ultra)sensitivity. We also estimated the area between the up and down branches of the response curves (*h*) to quantify hysteresis. We found that both n_H_ and *h* increased monotonically with G when either VIP cells (Fig. 4a) or PV cells (Fig. 4b) were stimulated. Within a limited range of *G*, the Hill coefficient (nH) increased monotonically for the transition from SST to VIP or SST to PV states in both layers, while hysteresis was absent (*h*=0). Hysteresis occurred at different values of *G* depending on the type of switch and layer and caused an abrupt increase in n_H_, which stabilized with large values of *G* and gave rise to a virtually binary state transition. Closer study of the bifurcation diagrams showed marked differences across switches and layers. As we increased *G*, the sensitivity increased rapidly for the VIP to SST switch in both layers. By contrast, a rapid increase of sensitivity for the PV to SST switch occurred at higher values of *G* in the superficial layer and even higher *G* values in the deep layer (Fig. 4c), as compared to the VIP to SST switch. Likewise, hysteresis onset increased from the VIP switches to the PV switch in the superficial and deep layer (Fig. 4d). Note that the PV switch in the deep layer showed an early increase of *h* due to propagated hysteresis effect from the superficial layer (*G* ~ 500, Supplementary Fig. 5c) and showed its own hysteresis increase later.

**Figure 4.**
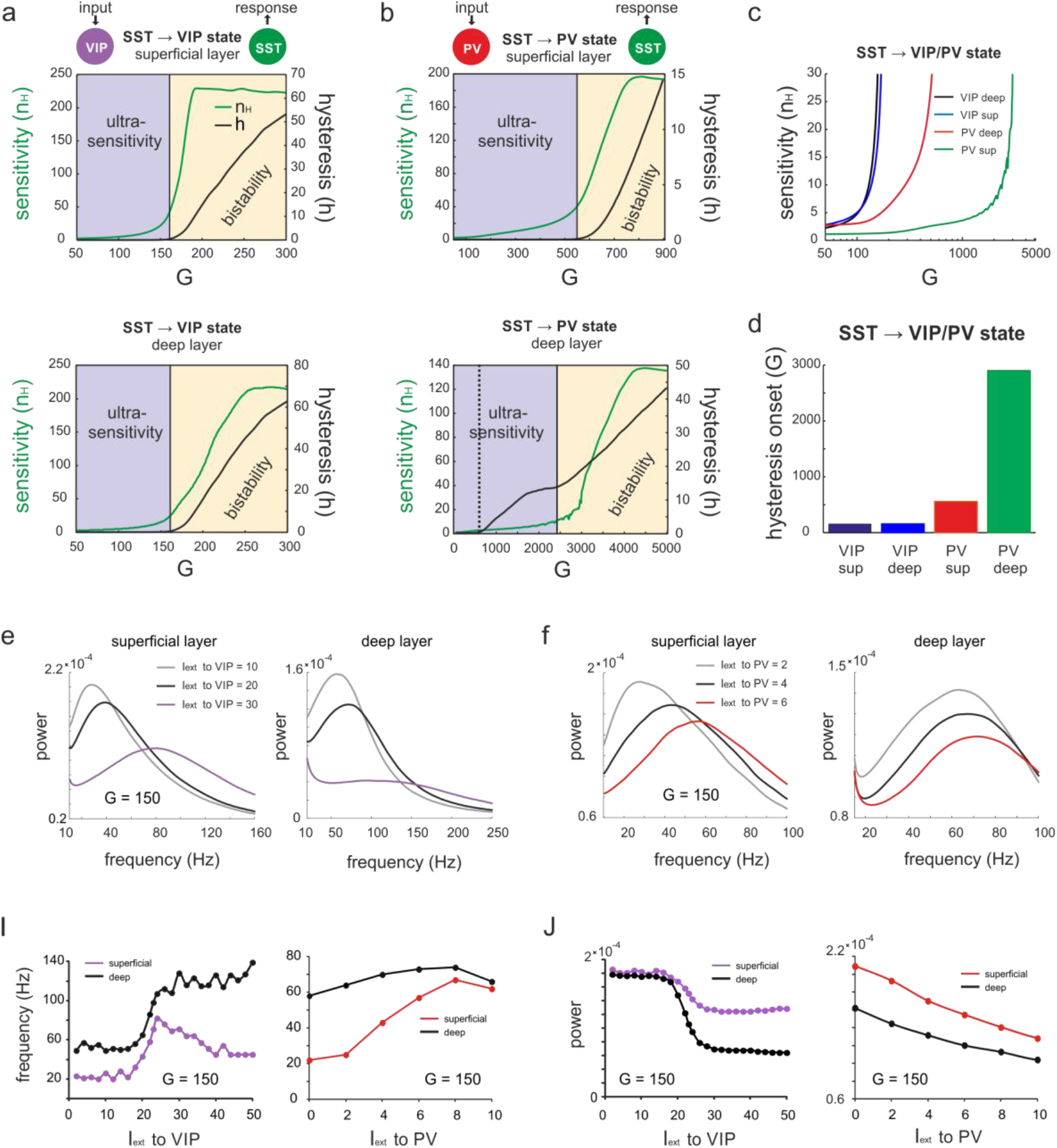
Bifurcation diagrams and spectral properties of two microcircuit switches. **a)** Bifurcation diagram of the superficial (top) and deep layer (bottom) depicting sensitivity, as measured by the Hill coefficient (N_H_), and hysteresis area (h) of SST cell responses to VIP cell input as a function of G. The shaded areas show a purely ultrasensitive and a bistable region, where both ultrasensitivity and hysteresis are present. **b)** Same as in a) for input to PV cells. Bottom dotted line: Onset of hysteresis in the superficial layer (top) is also seen in the deep layer (bottom) **c)** Sensitivity of SST responses to VIP or PV input in superficial and deep layers as a function of *G*. **d)** *G*-values of hysteresis onset in SST responses in both layers after input to VIP and PV cells. **e)** LFP power spectra of superficial and deep layers with three different inputs to VIP cells and a constant drive I_ext_ = 5 to SST cells. **f)** Same as in e) for input to PV cells. **i-j)** Peak frequency i) and power j) in superficial and deep layers as a function of VIP and PV input.

Subsequently, we studied the outcome of the switching dynamics on oscillation frequency and power of the LFP. Both input to VIP and PV cells strongly increased the frequency of the dominant oscillation in superficial and deep layers, whereas the power of the oscillation peak generally decreased (Figs 4e-j). This is expected from a transition from an SST dominated state with low frequency to a high oscillation state in which the frequency is imposed by dominating PV activity (see above). Note that frequency and power show sigmoidal jumps similar to the rate transitions above (Figs. 2i-j, VIP input).

Taken together these results suggest that the microcircuit can operate in two different states: An inhibited state, where SST neurons dominate and a disinhibited state with prevailing PV activity. Two mutually inhibitory circuit motifs provide two switches with different sensitivity which toggle the network between both states, characterized by different PYR rates and oscillation frequencies, while maintaining sufficient inhibition to putatively prevent runaway excitation.

### Lateral inhibition switches circuit to the SST state

Next, we studied switching dynamics in the microcircuit in the opposite direction, i.e. a transition from PV/VIP toward SST governed activity. To this end we applied input to SST cells in both layers (Fig. 5a), mimicking the effect of lateral inhibition during surround suppression in the visual cortex, which was experimentally found to be mediated by horizontal pyramidal cell input from distant microcircuits to SST neurons^45,33^. Driving SST cells resulted in a monotonic decay of activity of all other cell types in both layers and for different values of *G* (Fig. 5b), in line with several experimental studies^24,45,46^. Stimulation of SST cells also reduced the frequency of oscillation (Figs. 5c-e). However, as we increased the input to SST cells the power of oscillations initially increased and subsequently decreased once PV cells were strongly suppressed. This finding replicates experimental findings, in which stronger surround suppression is followed by a sudden transition from high frequency (gamma range) to lower frequency oscillations (high beta, low gamma range) and a concomitant increase in oscillatory power in mice^24,33^ and monkeys^47^.

**Figure 5.**
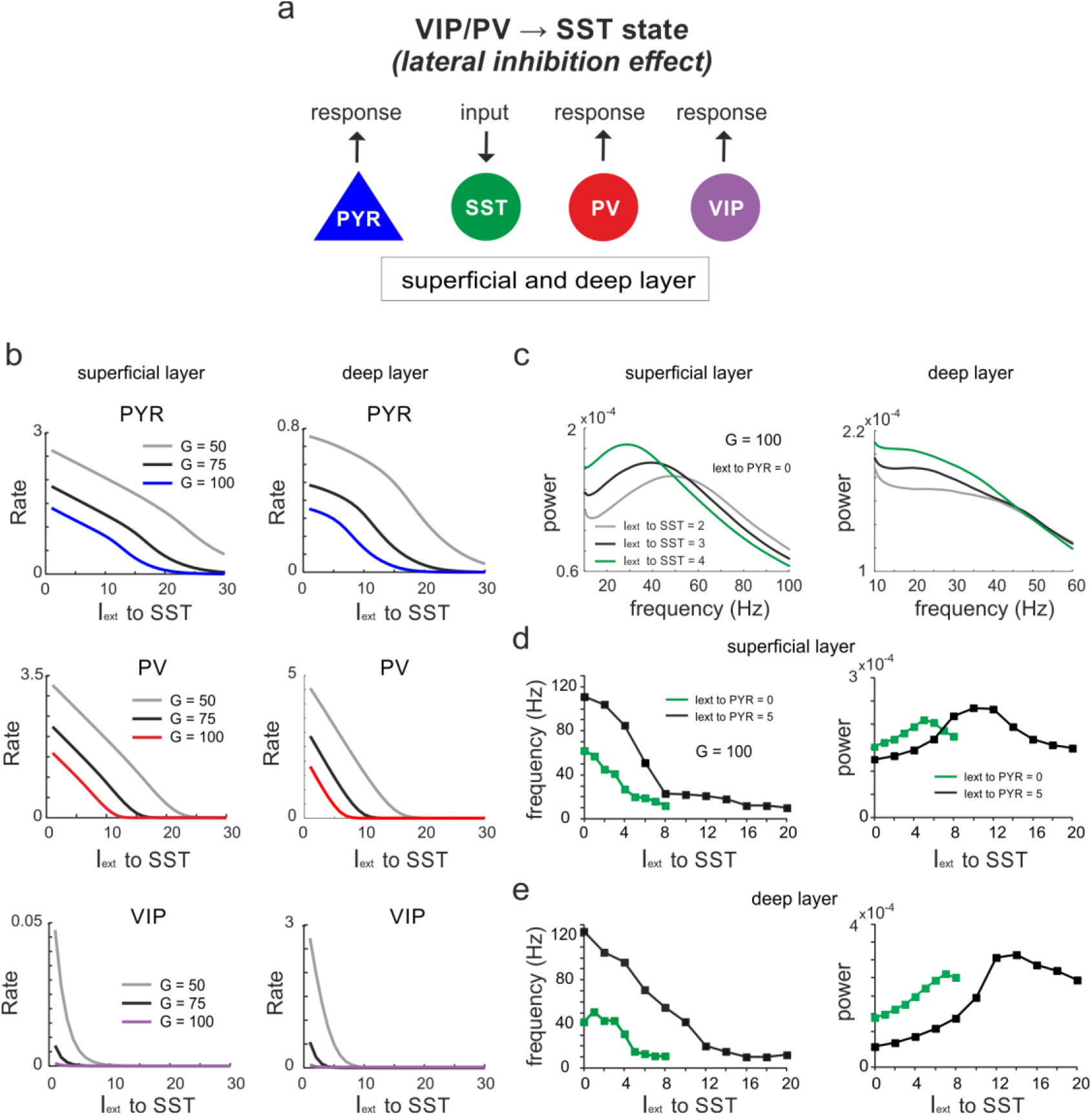
Lateral inhibition switches dynamics from VIP/PV to SST dominated states. **a)** Schematic showing input to SST cells in both layers and measuring response in all other cell types. **b)** Response curves of PYR, PV and VIP cells as a function of SST drive for superficial and deep layers, and three different values of *G*. **c)** Power spectra of superficial and deep layers for different levels of SST input. **d)** Peak frequency and power as a function of SST input for two levels of PYR drive. **e)** Same as in d) for the deep layer.

### Input to Pyramidal cell favors SST activity

Next, we measured the response of SST cells when PYR cells were stimulated in both layers (Fig. 6a-c, Supplementary Fig. 7a). We studied the response of SST cells in three different scenarios. In the first scenario we stimulated PYR cell and VIP and PV cells received no external input. In this scenario, PYR input strongly increased SST activity (Fig. 6.a) (and to a lesser extent PV rates, see Supplementary Fig. 7b) in both layers, while VIP cells were suppressed (Supplementary Fig. 7b). Stimulation of PYR cells also increased the population oscillation frequency, whereas oscillation power decreased (Supplementary Fig. 7c). In a second scenario we stimulated the PYR cells while VIP cells received a constant external input. In this scenario, sufficiently high PYR input and *G*-value caused a jump back to higher SST activity in both layers (Fig. 6b) with a sudden suppression of PV and VIP activity (Supplementary Figs. 8a-b). In contrast to the PYR cells, stimulation of VIP cells affected the oscillation and their power in a nonmonotonic fashion: the oscillation frequency increased initially, but dropped (superficial layer) or saturated (deep layer) at the transition to the SST state, while oscillatory power declined and suddenly increased with the switch to SST activity (Supplementary Fig. 8c). Finally, we also stimulated PYR cells while PV cells (instead of VIP cells) received a constant external input. In this scenario we obtained results similar to those obtained in the second scenario (Fig. 6c, Supplementary Figs. 9a-c). These findings are consistent with recent optogenetic experiments in which strong PYR drive was associated with high SST activity and comparatively low PV rates^48^.

**Figure 6.**
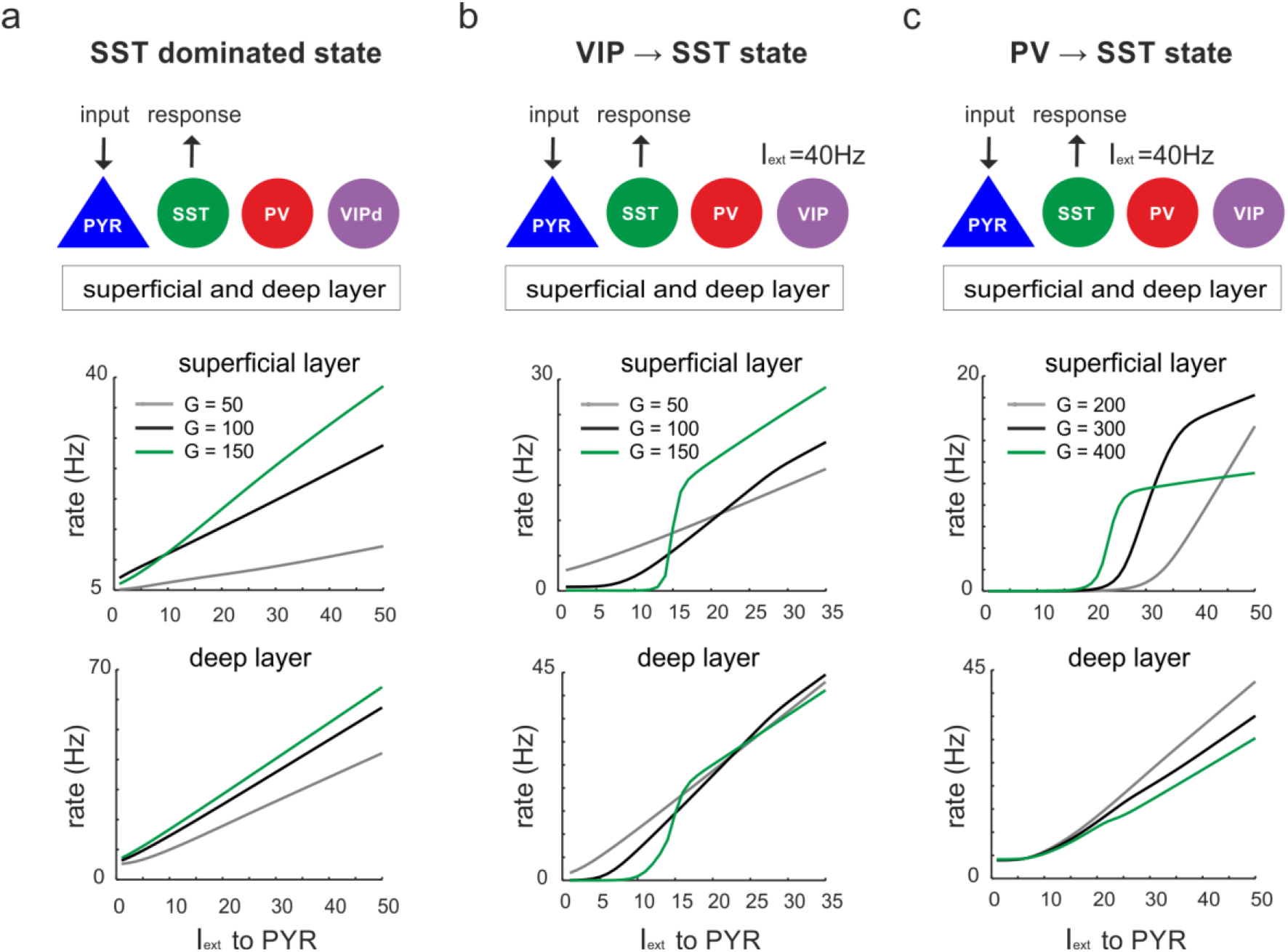
Input to PYR cells poises the microcircuit to SST dominated dynamics. **a)** Schematic of PYR cell input (top) and response of SST cells to increasing PYR drive in both layers for three values of *G* (top and bottom). **b)** Same as in a) with VIP cells being driven with a constant current I_ext_ = 40. **c)** Same as in a) with constant input I_ext_ = 40 to PV cells.

### State changes propagate between superficial and deep layers

Next, we addressed the question, whether a state transition triggered in only one layer propagates to the other layer across translaminar connectivity. To this end, we induced the same state changes as studied above (Figs. 3–5) by applying current to specific inhibitory neurons in only the superficial or deep layer and measured the response of pyramidal cells in the opposite layer (Fig.7). In addition, we removed translaminar connections to test their role in the state propagation (only connections with an impact are shown). When the circuit displayed high SST activity, input to VIP cells in the superficial layer showed a disinhibitory increase in the deep layer, which was mainly due to a translaminar reduction of SST activity, rather than a direct drive from superficial to deep PYR cells (Fig. 7a). We obtained a similar result in the superficial layer after driving the deep layer VIP cells (Fig. 7b). Likewise, input to PV cells in the superficial layer caused disinhibition in the deep layer within a certain input range which was abolished by removing translaminar SST connections (Fig. 7c). Notably, translaminar PV connections reduce the disinhibition effect as their removal strongly augmented PYR activity. The same effects were found in the superficial layer after PV input to the deep layer (Fig. 7d). When we set the circuit to a VIP dominated state (I_ext_ to VIP in both layers = 40) and applied current to SST cells in the superficial layer, PYR cell activity was suppressed in the deep layer due to a direct translaminar SST connection (Fig. 7e). The same results held true for SST input to the deep layer (Fig. 7f). Finally, a similar suppressive effect on PYR cell rates that propagated to the other layer after input to SST cells was found in the presence of the PV dominated state (I_ext_ to PV in both layers = 40), again due to the translaminar SST connections. (Fig. 7g-h). In summary, these results demonstrate that transitions between disinhibited and inhibited states triggered in one layer can propagate to the opposite layer and this interlayer interaction is primarily governed by translaminar SST connections.

**Figure 7.**
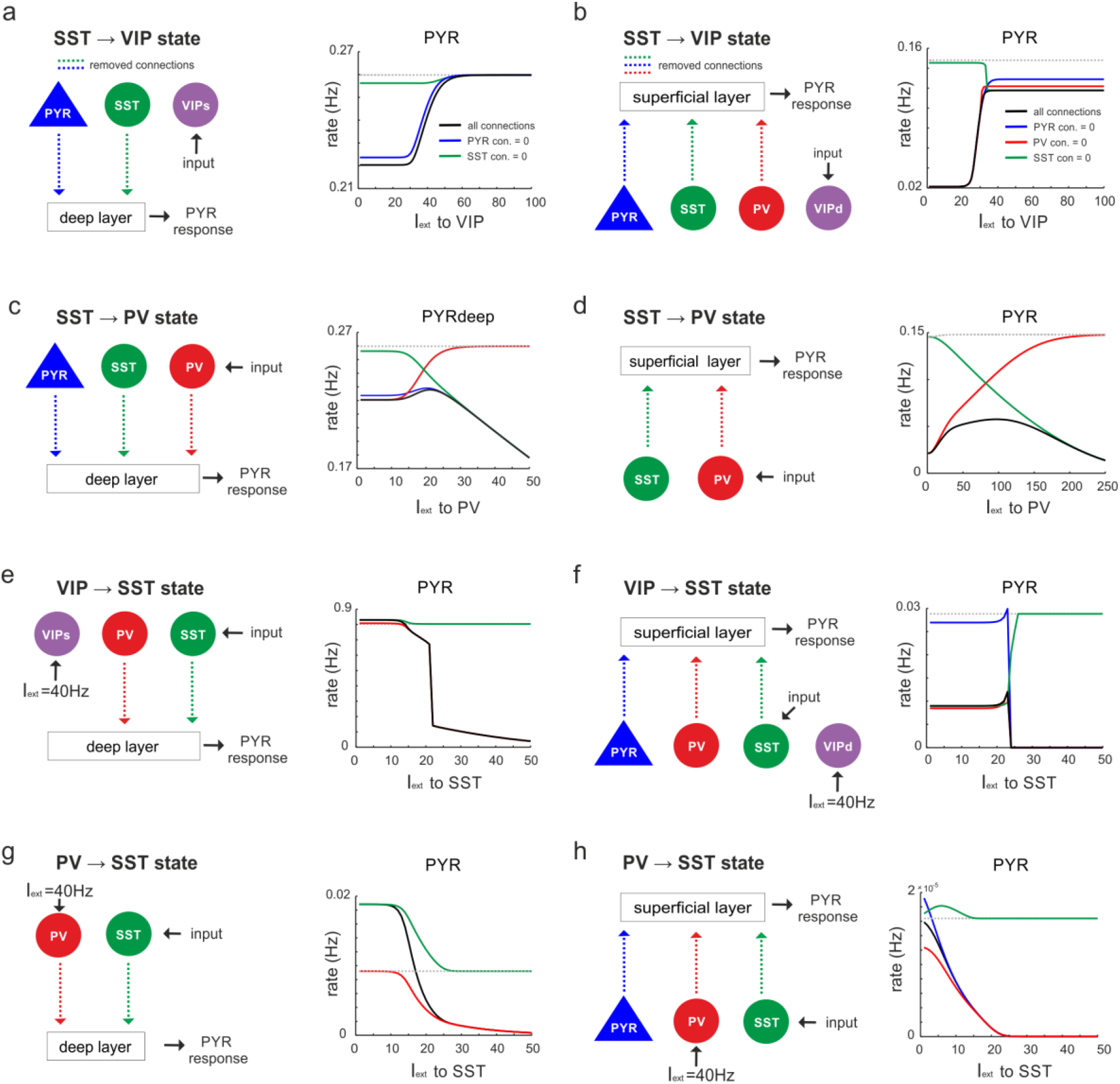
State changes propagate across layers. **a)** Input was only given to VIP cells targeting the superficial layer and the PYR response was measured in the deep layer after all connection from superficial PYR or SST cells to deep layer cells (dashed lines) were severed (left). The PYR response is shown for the case with intact connections (gray dashed line), all indicated connections cut (black solid line) or individual connections removed (colored lines). **b-h)** Same as in **a)** for input to different cells and different layers. Only connections were removed that had a visible influence on the PYR response as compared to the case where all connections were left intact.

### Recurrent inhibitory connectivity differentiates inhibited from disinhibited states

The connectivity between different interneuron types and pyramidal cells allows the microcircuit to switch between a disinhibited state with high PYR cell firing and an inhibited state with reduced PYR activity, as we show above. However, for a deeper understanding of this switch one important question remains: Why are PYR cells more inhibited in the SST dominated regime as compared to PV governed regime? To this end we examined the mouse V1 connectivity matrix and found that there is a strong asymmetry between recurrent connections among PV cells and SST cells. While connections between PV cells in each layer are the strongest within the entire matrix, recurrent connections among SST cells are entirely absent (Figure 8a, Supplementary Table 4). Thus, we hypothesized that the strong self-inhibition of PV cells effectively reduces their inhibitory effect on PYR cells while SST cells can elicit strong inhibition on PYR cells due to the absence of self-inhibition. If this was true, exchanging the recurrent connections, i.e. removing PV self-connections and add them to SST cells may invert their role in microcircuit state switching (Fig. 8a). In simulations with the inverted connectivity scheme we found that input to SST cells disinhibited PYR cells through release from PV and VIP suppression in both layers (Fig. 8b, Supplementary Figs. 10, 11a). Likewise, driving VIP or PV neurons was followed by inhibition of all the other cell types, similar to the effect of SST input in the original case (Figs. 8c-d, Supplementary Figs. 10, 11b-c). A notable exception is disinhibition in deep layer after PV input (Supplementary Fig. 11c). These results indeed suggest that inhibitory state dependent alterations in excitation-inhibition balance, which are at the core of the switching properties of the microcircuit, are due to large differences in PV and SST self-connectivity (Fig. 8e).

**Figure 8.**
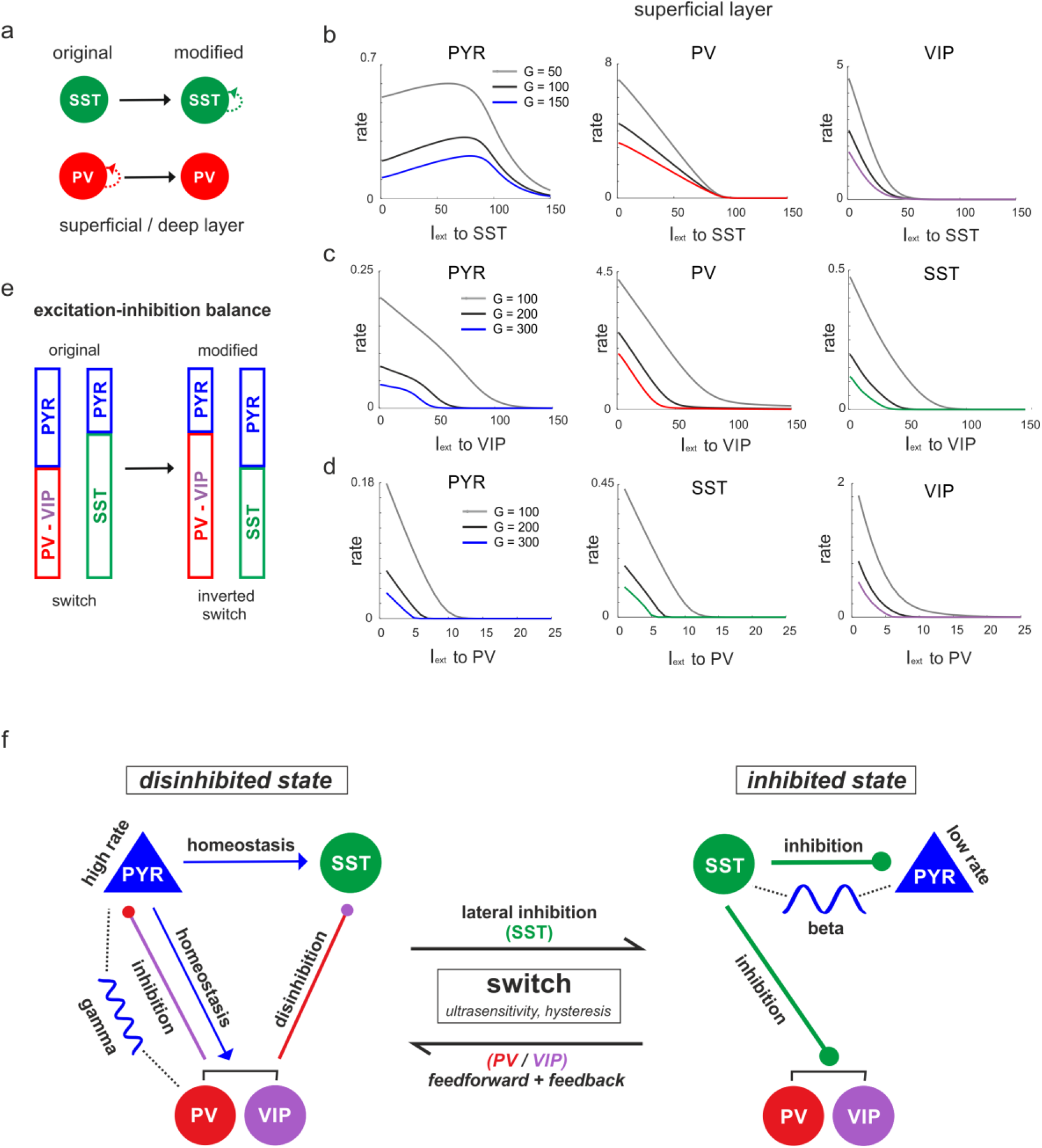
The microcircuit acts as a homeostatic switch between different levels of excitation and inhibition balance. **a)** Schematic showing the typical pattern of strong recurrent connections between PV cells and their absence in SST cells in superficial and deep layers (left). The pattern was inverted in the modified connectivity matrix (right). **b)** Response of PYR, PV and VIP cells to SST input using the modified connectivity matrix and three different values of *G*. Note the increase in PYR activity with moderate SST input. **c)** Same as in b) for input to VIP cells and response of PYR, PV and SST cells. **d)** Same as in a) for response of PYR, SST and VIP cells to PV input. **e)** Schematic showing how the excitation-inhibition balance changes between PV/VIP and SST dominated states using the original (left) or modified connectivity matrix, as shown in a). Excitation is mediated by PYR cells and inhibition by VIP/PV or SST cells. **f)** Summary diagram displaying the principles of the homeostatic switch implemented in the connectivity matrix of the studied microcircuit.

## Discussion

In this study, we showed that the mouse V1 microcircuit is endowed with two switch like mechanisms that can toggle the pyramidal cells (output of the microcircuit) between high (disinhibited) and low activity states (inhibited) across superficial and deep layers (Fig. 8f). The underlying switching mechanics are realized by the interactions among the three interneuron types (PV, SST, VIP), which compete for inhibitory influence on pyramidal cells. In the inhibited state, SST cells dominate inhibition which serve as ‘master regulators’ by strongly connecting and inhibiting activity of pyramidal cells and other interneurons in the circuit^24,34,49–51^. In the disinhibited state, excitation is mainly balanced by PV cells^52^ and to a lesser extent by VIP neurons^53^, whereas SST activity is reduced.

### Difference between inhibition exerted by PV and SST neurons

While disinhibition through SST suppression was previously shown experimentally and theoretically ^18,19,27,54^, the question remains why SST cells provide more inhibition than PV or VIP cells, even though the weights of the PV to PYR connections by far outweigh SST to PYR connections (Supplementary Table 4). Similar to previous simulations of simplified neuronal networks^27^, we found that a key to this inhibitory asymmetry lies in the degree to which PV and SST cells are connected among themselves. While SST cells lack mutual connectivity, PV cells have strong mutual connectivity (that may also be reinforced by gap junctions^6,7,55^). In our rate model, self-inhibition of PV cells reduced their impact on PYR cells and enhanced PYR rates. We note a similar effect was found in spiking network models, where PV interaction enhances PYR rate and synchrony^56^. Accordingly, we found that exchanging self-connections of PV and SST neurons inverted their role in mediating the inhibited or disinhibited state.

### Origin of switching dynamics

The switching mechanism emerges as a consequence of two mutual inhibitory connection motifs i.e. from SST to PV and VIP, and PV / VIP to SST neurons, in both layers. Thus, driving SST cells switches the microcircuit state to the ‘inhibited state’ by suppressing PV and VIP neurons as well as PYR neurons. Conversely input to VIP or PV cells disinhibits the circuit by decreasing SST cells activity. The nature of this switch depended on the mutual inhibitory connectivity strength, rendering the state transition purely ultrasensitive with sigmoidal input-output curves for weaker weights or bistable with hysteresis and memory for stronger weights, consistent with recent observations made in simplified inhibitory neuronal networks^27^. Switching based on double negative feedback has been conserved during evolution in many biological systems^43,44,57^ and we found that it may also be a hallmark of cortical microcircuits. Notably, the two ways to control the switch differ in their sensitivity, with VIP cells requiring considerably less input to control SST activity and flip the circuit to the disinhibited state in both layers than PV cells. Thus, PV cells seem to be more specialized in keeping the excitation-inhibition balance at sufficiently high levels, whereas VIP cells play more a switching role, even though both cell types can assume both functions.

The switching circuitry of the microcircuit can not only be activated by simultaneous input to superficial and deep layers, but each layer can transmit its state change to the other through translaminar connectivity, and effectively synchronize inhibited or disinhibited states across the whole microcircuit. These results suggest that the two-layer microcircuit as a whole can act as a switch, whose state may be used to guide activity flow of pyramidal cells in the feedforward (via L2/3) and deep excitatory cells in the feedback direction (L5)^58,59^. However, it is conceivable that differential input to superficial and deep layers could place both in different switch states.

### Bistable dynamics

The presence of bistability based on mutual inhibitory connection motifs also provides an alternative to the dominant view that persistent transitions between low and high firing rate states across the brain^60^, which underlie working memory required for attention^61^, consciousness^62–64^ and language processing^65–67^, are primarily based on local^68^ or inter-areal^62,64^ recurrent connectivity between excitatory neurons. In our study, bistability is controlled by properties of local inhibitory connectivity, opening up the possibility that anatomical heterogeneity and gradients across the brain may reflect the presence of bistable switches in some brain areas and ultrasensitive switches without memory in other regions^69–71^.

### Oscillations in the microcircuit

Two negative feedback loops endow both deep and superficial layers with an intrinsic oscillation. Each loop has a different frequency owing to differences in synaptic time constants of PV and SST cells^72^. Consistent with experiments and models, PV cells drive high frequencies (‘gamma range’), whereas SST cells prefer lower frequencies (‘beta range’) ^22,24,29^. Accordingly, during the inhibited switch state the dominating SST cells impose a slower frequency on PYR cells as compared to the PV dominated disinhibition state with high frequencies. Animal experiments have revealed that the power of high frequency oscillations was stronger in the superficial layer, while slower oscillation had more power in the deep layers^32,39–41^. Based on our results we argue that emergence of such differences in oscillations in deep and superficial layers crucially depended on specific anatomical connections (see Fig 2a). In intermediate switch states, the final frequency was established by the relative contribution of PV and SST cells, which provides a potential explanation for a longstanding question of what determines the oscillation frequency in neuronal circuits^73,74^.

Thus, our results also suggest an alternative taxonomy for oscillations, dividing them into competing SST (low frequency) and PV driven rhythms (high frequency). Notably, during the transition between (PV or SST dominated) switch states the oscillation frequency and power behave asymmetrically i.e. an increase in frequency is accompanied by a decrease in power. However, when PV or SST cells are strongly suppressed or silenced, frequency and power change symmetrically. A decrease in frequency and increase in power is also found in recent experiments in monkeys and mice, in which small stimuli trigger a high frequency oscillation (~60 Hz) that is replaced by a lower frequency (~30 Hz) in the presence of larger stimuli^4,33,47^.

Anatomical studies show that this surround suppression effect is presumably mediated by lateral excitatory input from the surround to SST neurons in the center^45^. Our model suggests that the surround switches the microcircuit from a PV/VIP dominated state with a high frequency putatively mediated by stimulus triggered feedforward and feedback drive^75,76^ to a more inhibited state caused by enhanced lateral drive to SST cells. We note that experiments have also reported the inverse case with an increase in oscillation frequency and decrease in power after enhancing visual stimulus contrast^77^.

### Stability of the cortical microcircuit

The disinhibited state of the switch naturally entails the risk of runaway excitation. Our results provide evidence that the microcircuit contains supplementary homeostatic mechanisms that keep disinhibition within healthy boundaries. When the circuit is disinhibited and SST activity suppressed, strong drive to pyramidal cell can cause a sudden reversal of SST neurons to a high firing rate state and thereby restore a more inhibited state in the microcircuit. This phenomenon was reported experimentally^78^ and in a recent theoretical study^28^ showing elevated visually evoked SST rates, after prior suppression through VIP cells activated during locomotion.

### Predictions

We found that asymmetric changes in oscillation frequency and power are abolished with silencing of PV or SST cells, a result that can be experimentally tested. Moreover, our model predicts that driving PYR cells is followed by linear or non-linear responses in inhibitory cells depending on the dominance of SST or PV/VIP cells, respectively. Finally, experiments could test whether inhibition or disinhibition mediated by driving specific inhibitory cells in one layer propagates to the other layer, as our simulations showed.

In conclusion, the function that emerges from our computational study of the microcircuit is a homeostatic switch that toggles pyramidal cells, the principal output neurons of the circuit, between an inhibited and disinhibited state. The switching dynamics is orchestrated by an array of inhibitory neurons, each performing a specific task in the switch mechanics. Feedforward, feedback and lateral input may change the position of the switch and regulate the flow of excitation to downstream microcircuits^10,15,79^. Our results also map different types of oscillations onto different interneuron types and link them with distinct switch states, which in the future may help to bring together rate and oscillation based experimental paradigms. They provide mechanistic insight into the long held notion that slow oscillations assume an inhibitory function, while fast oscillations serve information processing^79–83^.

## Methods

### Microcircuit architecture

In this study we develop a firing rate model of the visual cortical microcircuit. This model is based on pair-wise connectivity between major neuron types in superficial and deep layers of the neocortex^34^. Using octuple whole-cell recordings, Jiang et al.^34^ exhaustively mapped out the connectivity (EPSP or IPSP strength and connection probability) between a large number of morphologically defined neurons (interneurons and pyramidal cells) within and between layers 1, 2/3 (superficial) and 5 (deep) of the primary visual cortex of the mouse, while layer 4 was not included. In addition, Jiang et al. ^34^ also quantified the relative prevalence of each cell type and reported synaptic plasticity properties of specific connections. Finally, based on genetic makeup different interneurons were labeled as PV^+^ cells (layer 2/3: basket cell and chandelier cells; layer 5: shrub cells and horizontally-elongated cells), SST^+^ cells (layer 2/3 and 5: Martinotti cells) and VIP^+^ cells (only layer 2/3: bitufted cells and bipolar cells).

To convert the original connectivity data (see Supplementary Table 1 for the connection probability matrix and Supplementary Table 2 for the weight (EPSP/IPSP) matrix) into a format suited for a computational model, we performed several manipulations. First, restricting our analysis to pyramidal cells and three major interneurons types (PV, SST and VIP), we created a single connectivity matrix (C) by element-wise multiplication of EPSP/IPSP strength (i.e. mean amplitude) and connection probability matrices. To generate a single class of PV cells in each layer, we added the weights of basket cells and chandelier cells in the superficial layer and the weights of basket cells, shrub cells and horizontally-elongated cells in the deep layer. After inspection of the connectivity matrix, we noticed that out of the two existing VIP cell types, one VIP cell type preferentially targeted other neurons in L2/3 (bitufted cells), whereas the other only innervated L5 (bipolar cells). Based on this observation, in our model we divided VIP neurons into VIPsup from VIPdeep cell types, even though both types were anatomically located in the superficial layer.

Thus, we had four neuron types in both deep and superficial layers. Instead of modeling populations of neurons with *N* cells for each class, we modeled each cell population using a single firing rate-based neuron. However, we adapted the connectivity weights according to their relative prevalence which is computationally less expensive. To this end, we first followed the general rule that there are roughly five times more pyramidal neurons in a microcircuit than interneurons. Therefore, all inhibitory weights in the matrix were scaled down by a factor of 0.2. Next, we multiplied the weights of each interneuron type with its relative prevalence (Supplementary Table 3). This resulted in the corrected 8 x 8 connectivity matrix (Supplementary Table 4). The resulting microcircuit is schematically shown in Fig 1A.

### Population activity model

The dynamics of each neuron type, i.e. pyramidal cells and interneurons, was modeled using the coarse-grained firing rate-based model (Wilson-Cowan model^84^). The dynamics of the full microcircuit can be written in vector form as:

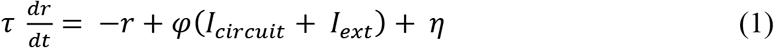

where *r* is the vector of rates of all 8 cell types, *τ* is the vector of population specific time constants (Supplementary Table 5), *η* reflects Gaussian noise with standard deviation *σ* = 0.01. The input-output firing rate transfer function (*φ*) of each neuron type was modeled as 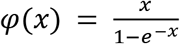 and is identical for all neuron types. The term *I_circuit_* denotes the input from other neuron populations across the entire microcircuit and is given by:

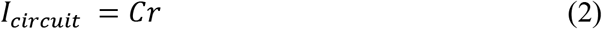

where C is the corrected connectivity matrix. *I_ext_* reflects external input to different neuron populations from bottom-up, top-down and lateral neuronal connections. Due to strong inhibition, PYR activity can be reduced to zero and even though the network is oscillating we may not observe those, as our readout of network oscillations is through PYR neurons. Therefore, it was necessary to inject some external input to PYR neurons I_ext_ to Pyr = 5 Hz (unless stated otherwise) in order to observe oscillations. The time constant *τ* was chosen for each cell type in accordance with the experimental and model literature, which attributed fast decay of PYR and PV activation and a longer decay for SST cells^24,72,85–87^. Because distances within the microcircuit are very short, conduction delays were not modeled explicitly. For the analysis of the rate response of a given cell class to input we performed a noise free (*σ* = 0) simulation of 2 seconds after which a steady state response was reached and the rate used for analysis (Figs. 3, 5, 6, 7). For Fig1c, 20 seconds were simulated, and the rate was averaged across the entire duration. To study oscillations we simulated 50 trials of 20 second each and averaged oscillation metrics across all trials. All simulations were performed with a time step of 0.1 ms.

### Control variables

To study the behavior of the microcircuit we varied several parameters. Foremost, we studied the dynamics of the microcircuit by systematically scaling the overall connectivity by a factor *G*. The scaling of the connectivity matrix can be loosely related to changing the absolute number of neurons in our circuit, similar to other studies ^35,36^. For the analysis we used the entire connectivity matrix without masking weak connections. The behavior of the circuit was studied by manipulating individual connections by removing or swapping them. Optogenetic silencing of individual cell types was simulated by setting specific columns of the matrix to zero i.e. we effectively removed all output of specific neuron types to the entire microcircuit (see Fig 2). To mimic stimulation of specific neuron types we injected direct current with varying amplitudes to the selected neurons type (e.g. see Figs. 3–6)

### Data analysis

#### Analysis of the LFP

We used the rate of pyramidal cells in each layer as a proxy for superficial and deep local field potentials (LFP) and its oscillatory behavior was investigated as a function of external drive. To analyze oscillation in the LFP we computed the power spectrum within a range of 1 to 250 Hz with the multitaper method implemented in the Chronux toolbox of Matlab (http://chronux.org/). The power spectrum was smoothed and normalized by the summed power of the computed frequency range. We then quantified visible oscillation peaks, excluding frequencies < 10 Hz, in terms of peak frequency and power.

### Measure of switching dynamics and hysteresis

Ultrasensitivity reflects the behavior of a system where small changes in input cuase large changes in output. Such behavior is observed in many natural systems such as biochemical reactions^43,88^. Ultrasensitivity can be quantified by fitting a sigmoidal curve to the input-output transfer function. Here we used the Hill equation^43^ to estimate the sensitivity of the output (y) to the input (x):

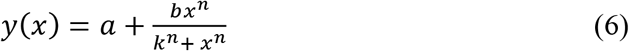

where is the intercept, b is the maximum, k is the half-maximum of the output *y* and *n* is the Hill exponent, which we used to quantify the response curves of SST cells to VIP and PV input (Figs 4a-c in the main text). If n = 1, the Hill curve is hyperbolic, whereas *n* > 1 indicates a sigmoidal shape with growing slope (i.e. sensitivity) as n increases.

Hysteresis in general describes the dependence of a system’s behavior on the past and implicates the presence of memory. As a consequence, system responses observed when input is steadily increased differ from responses to decreasing input. To test for hysteresis, we computed response curves of all cells (Fig. 3c-d, Supplementary Fig. 5) for ascending and descending VIP and PV input separately. For each input value (i), the rate response of all cells of the previous input value (i-1) was used to initialize the cell rates for the new input value. In the presence of hysteresis, the response curves for increasing and decreasing input do not collapse. Hysteresis (h) was then quantified as the summed difference of the ascending and descending rate curves of the SST cells in superficial and deep layers.

## Acknowledgments

G.H., H.S., T.R.K and G.D were funded by the German Research Council (DFG, No. KN 588/7 – 1) within the priority program Computational Connectomics (SPP 2041). G.D. was supported by the Spanish Research Project COBRAS PSI2016-75688-P (AEI/FEDER, EU), and by the Catalan AGAUR program 2017 SGR 1545. G.D. and G.H. received support from the European Union’s Horizon 2020 research and innovation program under Grant Agreement No. 720270 (Human Brain Project SGA1) and No. 785907 (HumanBrain Project SGA2). G.H. was funded by the grant CONSCBRAIN (n.661583) of the European Union’s Horizon 2020 research and innovation program under the Marie Skłodowska-Curie action. A.K. received support from the Swedish Research Council.

## Author contributions

G.H., A.K., H.S., T.R.K. and G.D. designed the study and interpreted the results. G.H. performed the simulations and analyzed the data. G.H. provided the first draft of the manuscript, which was reviewed and edited by the remaining authors.

---

**Supplementary Figure 1.**
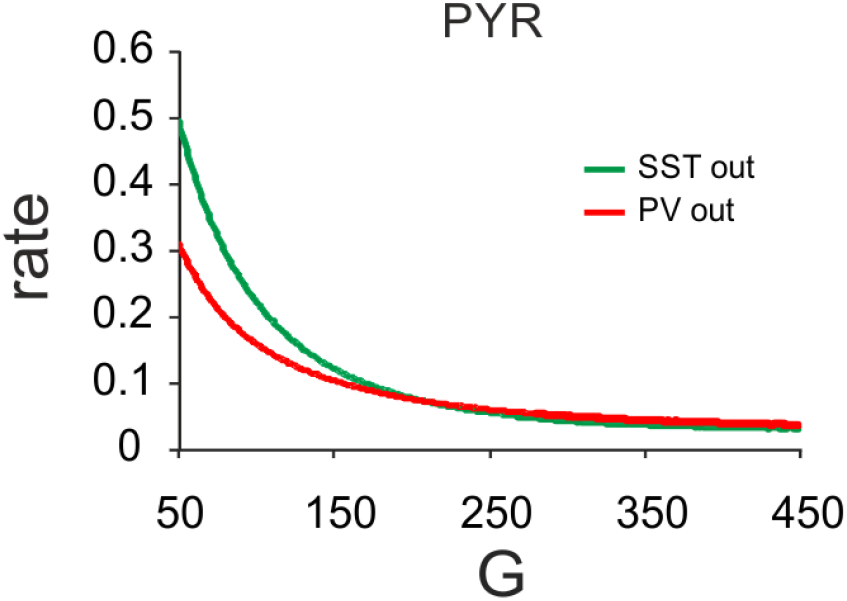
Comparison of the evolution of SST and PV rates with increasing *G*.

**Supplementary Figure 2.**
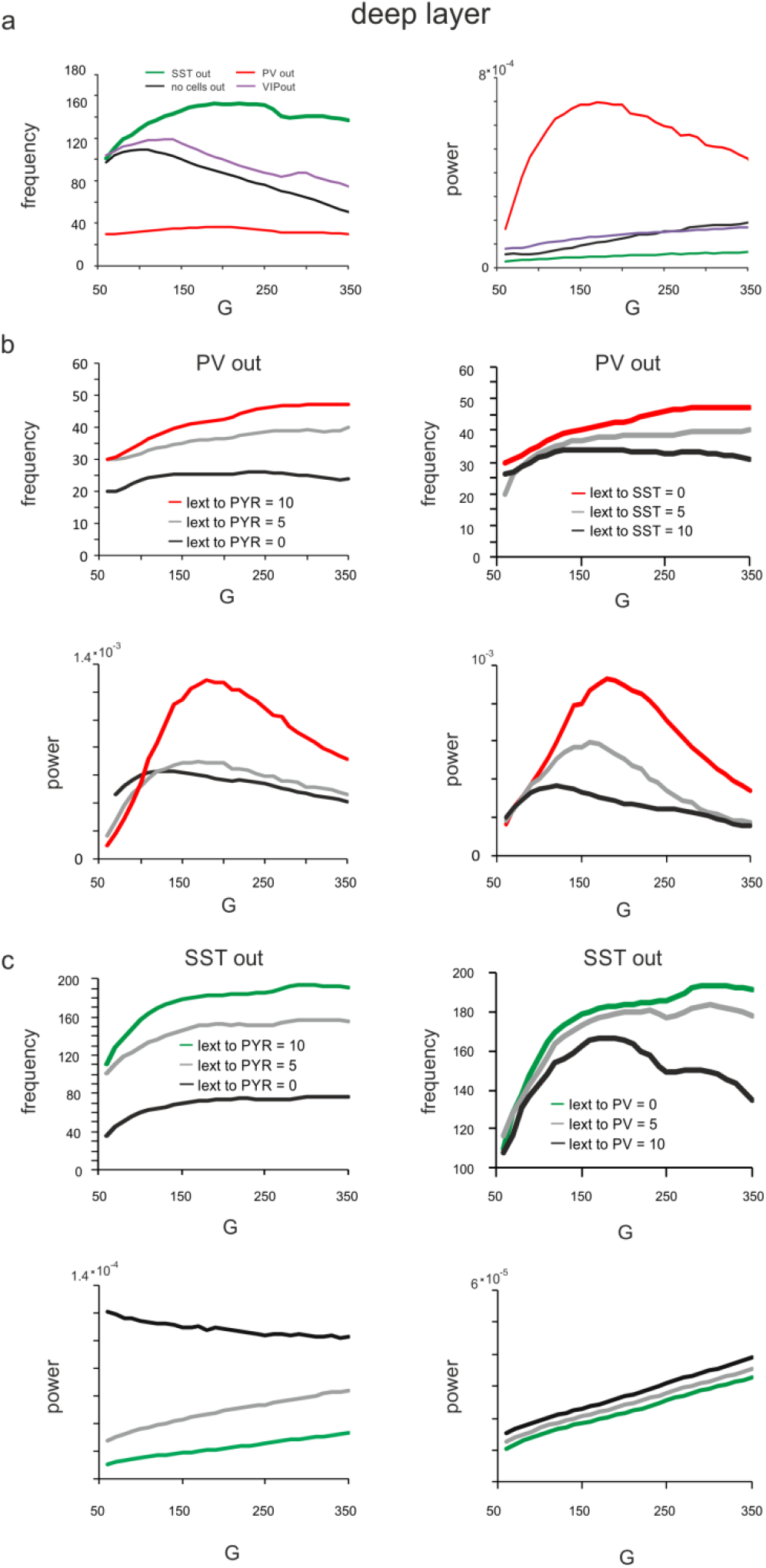
Effect of silencing specific cell types on rate and spectral properties in the deep layer. **a)** Peak LFP frequency (left) and power (right) as a function of *G* for different cell lesions in the deep layer. **b)** Peak frequency (top) and power (bottom) in the deep layer as a function of g after PV cell inactivation and different levels of input to PYR (left) or SST cells (right). **c)** Deep layer peak frequency (top) and power (bottom) after SST lesion as a function of g and varying input to PYR or PV cells.

**Supplementary Figure 3.**
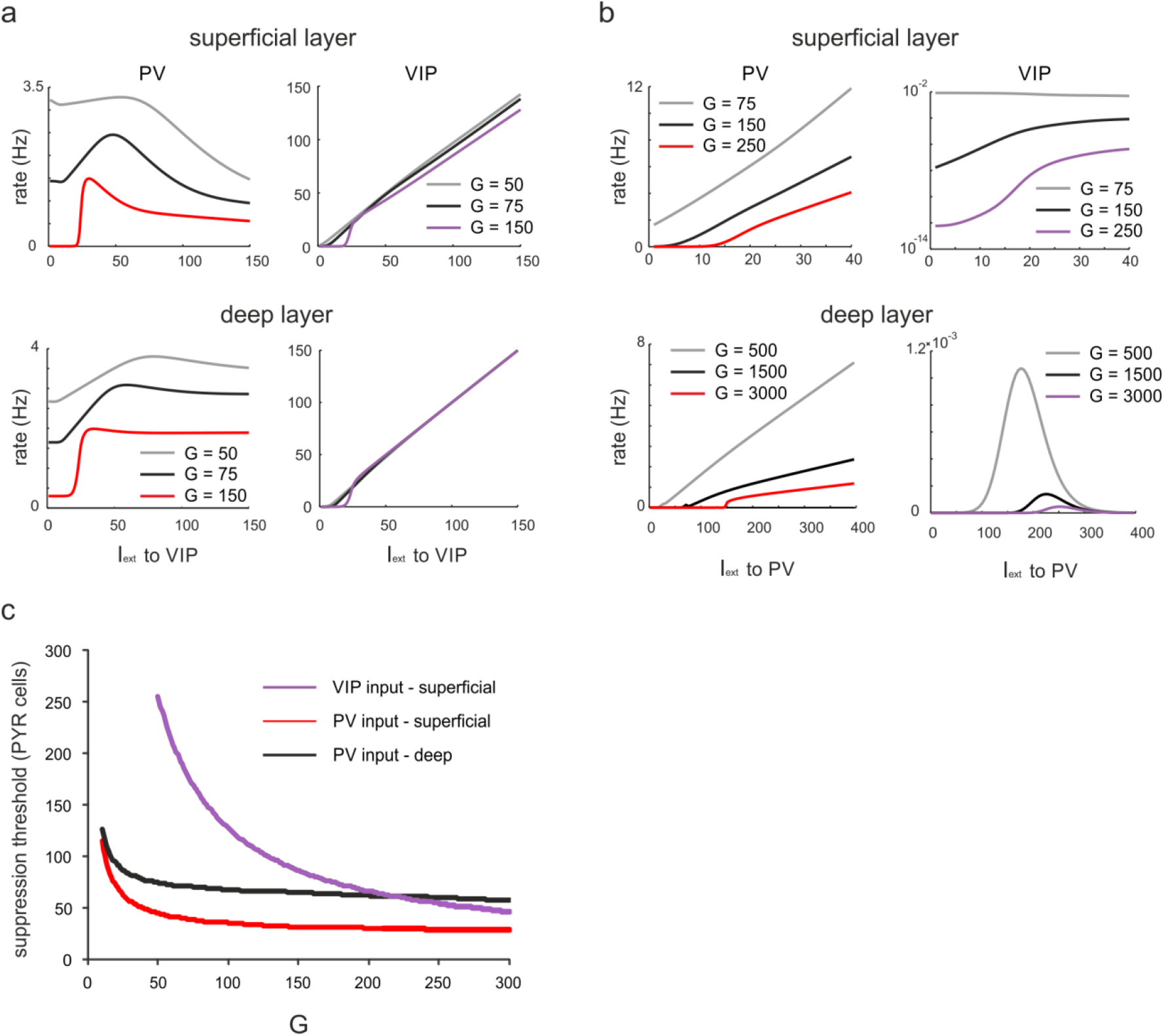
Ultrasensitivity in PV and VIP cells. **a)** Rate of PV and VIP cells in superficial and layers after input to superficial and deep VIP cells. SST cells were driven with a constant input I_ext_ = 5 to increase SST activity. Responses are shown for three different values of *G*. **b)** Same as in a) for simultaneous input to superficial and deep PV cells. **c)** Suppression threshold of PYR cells in different layers as function of *G*, when input was given to VIP or PV cells. The threshold was defined as the necessary input rate to PV/VIP cells, applied simultaneously to both layers, to suppress PYR rate to <0.001. Note that no threshold exists for PYR cells in the deep layer, as they are not targeted by VIP cells.

**Supplementary Figure 4.**
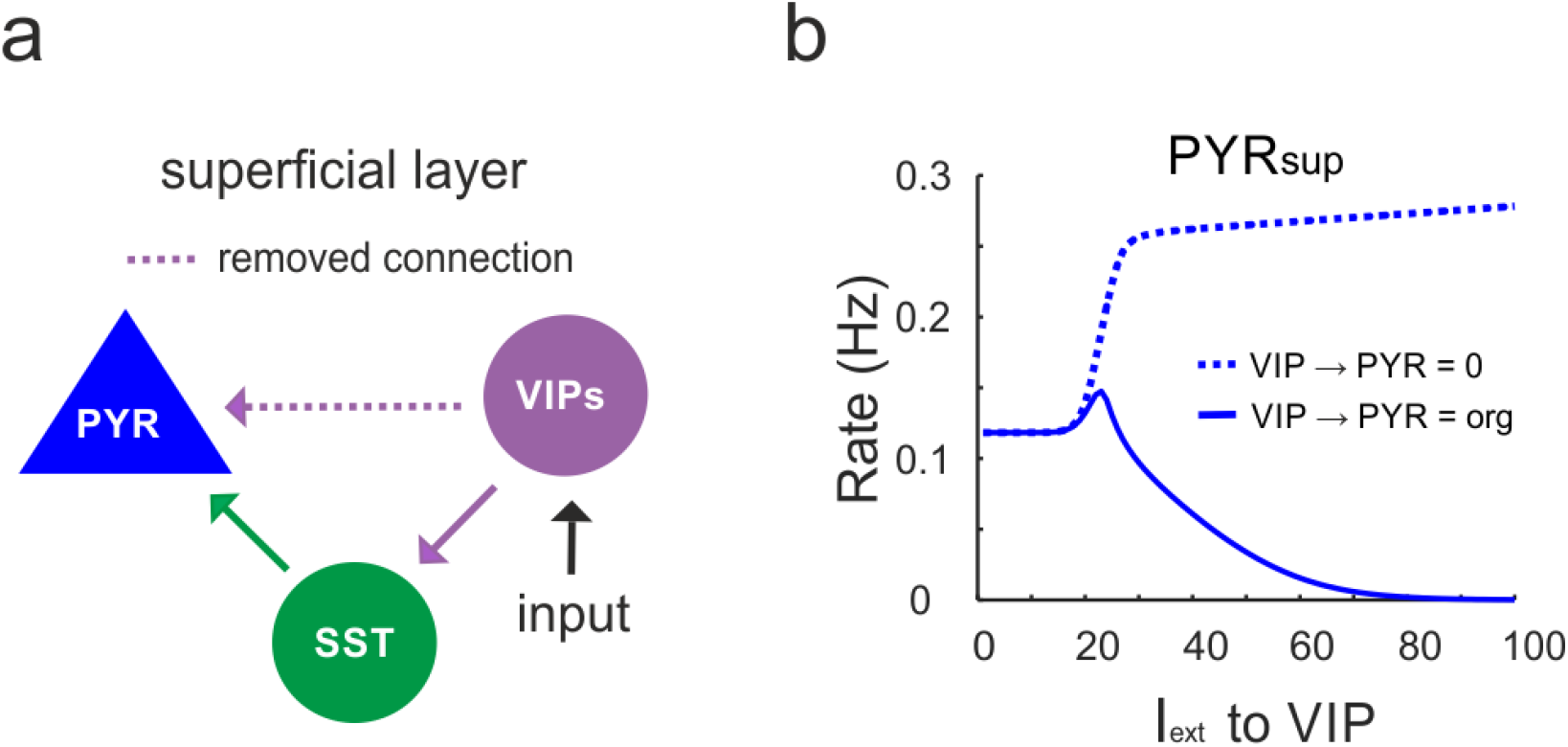
VIP cells only inhibit superficial PYR cells. **a)** Diagram in which the connection between superficial VIP cells and PYR cells was selectively removed. **b)** Comparison between original response of PYR cells to VIP input (solid line) and after removal of the connection, as shown in a).

**Supplementary Figure 5.**
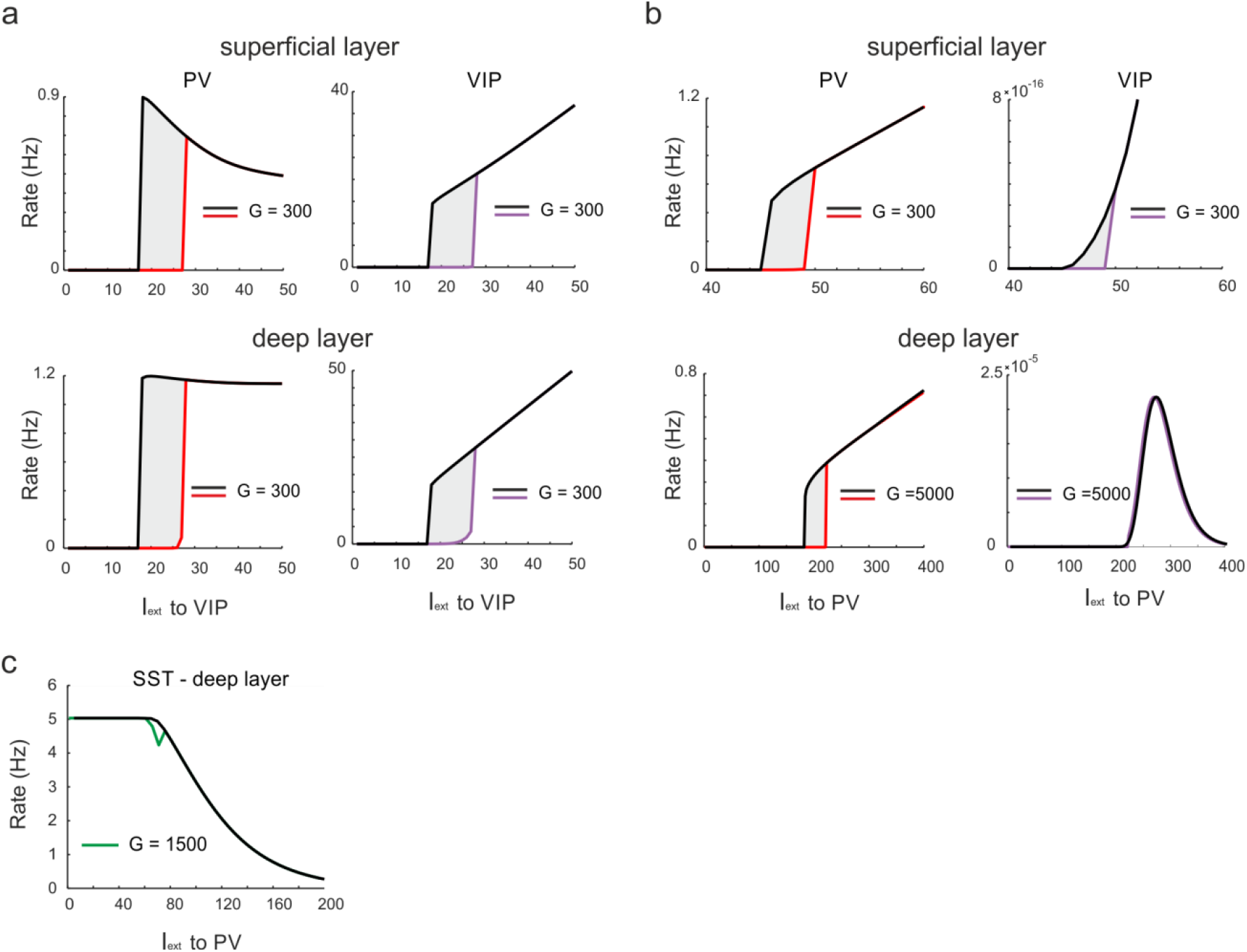
Hysteresis in PV and VIP cells. **a)** PV and VIP cell response to increasing (up branch) and decreasing input (down branch) to VIP cells for an exemplary value of g and both layers. The network displays hysteresis in each layer (shaded region h). **b)** Same as in a) for simultaneous input to PV cells in both layers. **c)** Response of SST cells in the deep layer to PV input and g = 1500.

**Supplementary Figure 6.**
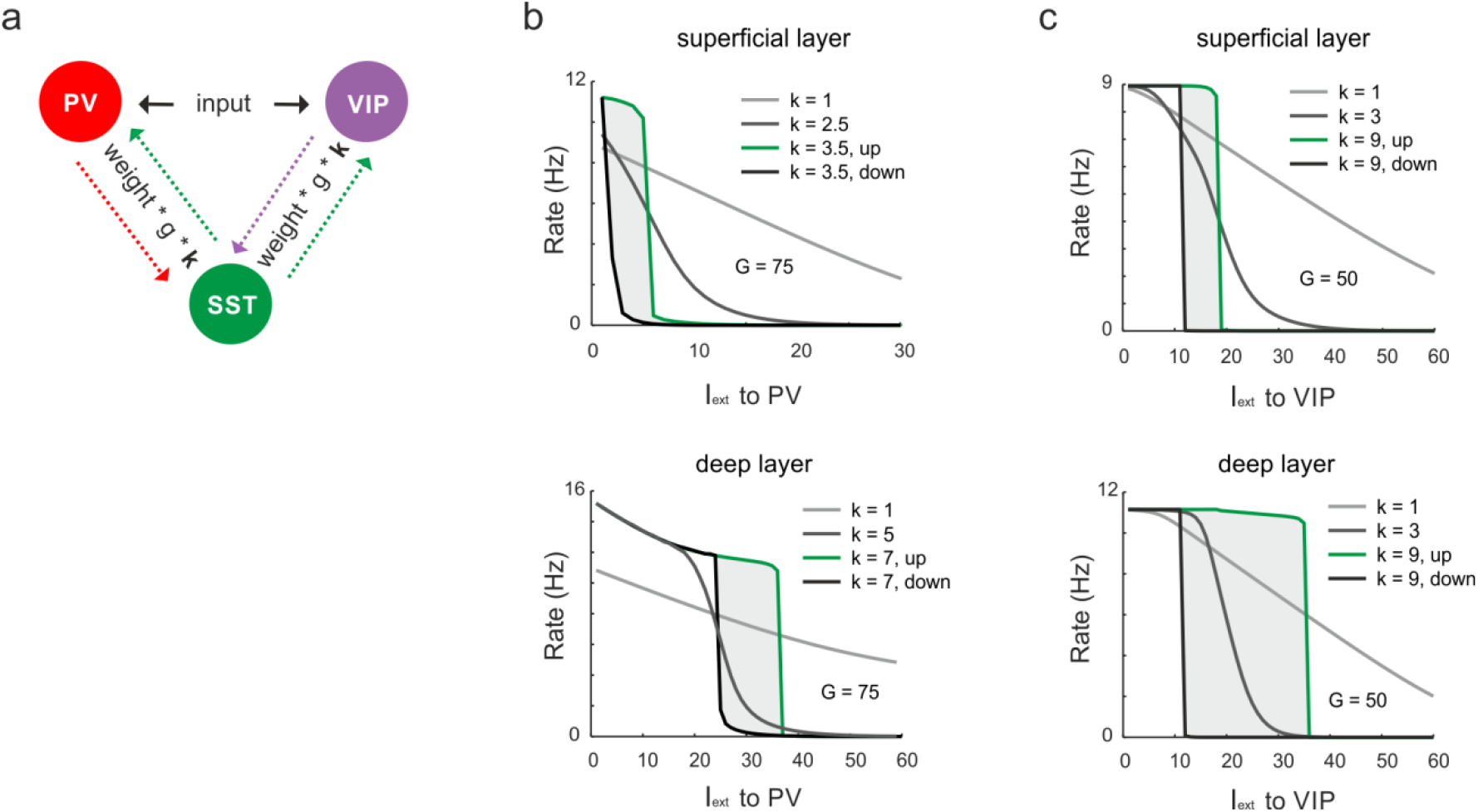
Mutual inhibitory connectivity weights control ultrasensitivity and hysteresis. **a)** Schematic showing the scaling of the connections between SST and VIP/PV cells with a constant k. **b)** Response of SST cells to PV input for different values of k and a constant value of *G*. **c)** Same as in (b) for input to VIP cells.

**Supplementary Figure 7.**
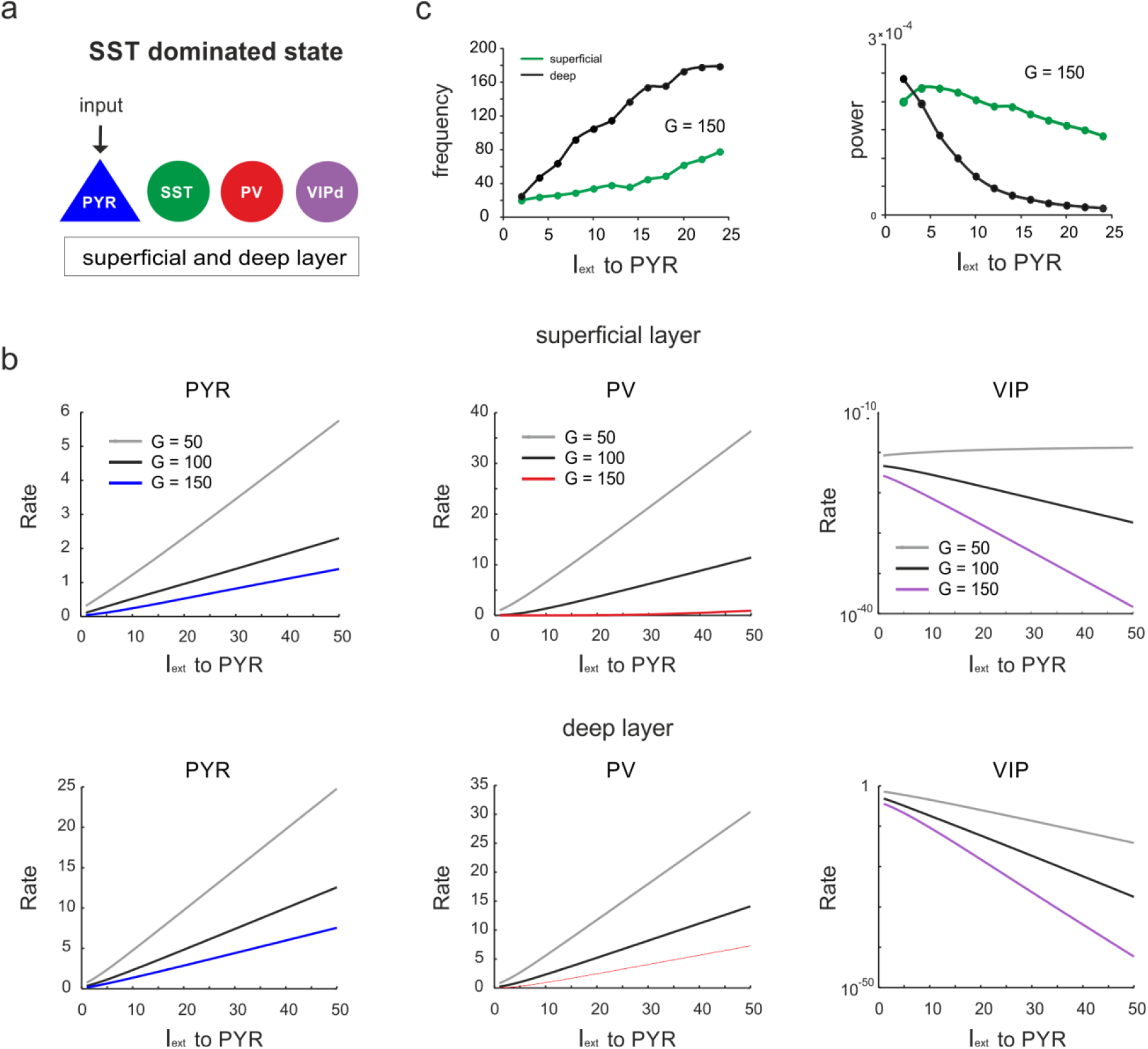
Response of different cell types to PYR cell input during the SST dominated state. **a)** Input was only provided to both superficial and deep PYR cells. **b)** Peak frequency and power in both layers as a function of PYR drive for an exemplary *G* value. **c)** Response of PYR, PV and VIP cells to PYR input for different values of *G* in superficial (top) and deep layers (bottom).

**Supplementary Figure 8.**
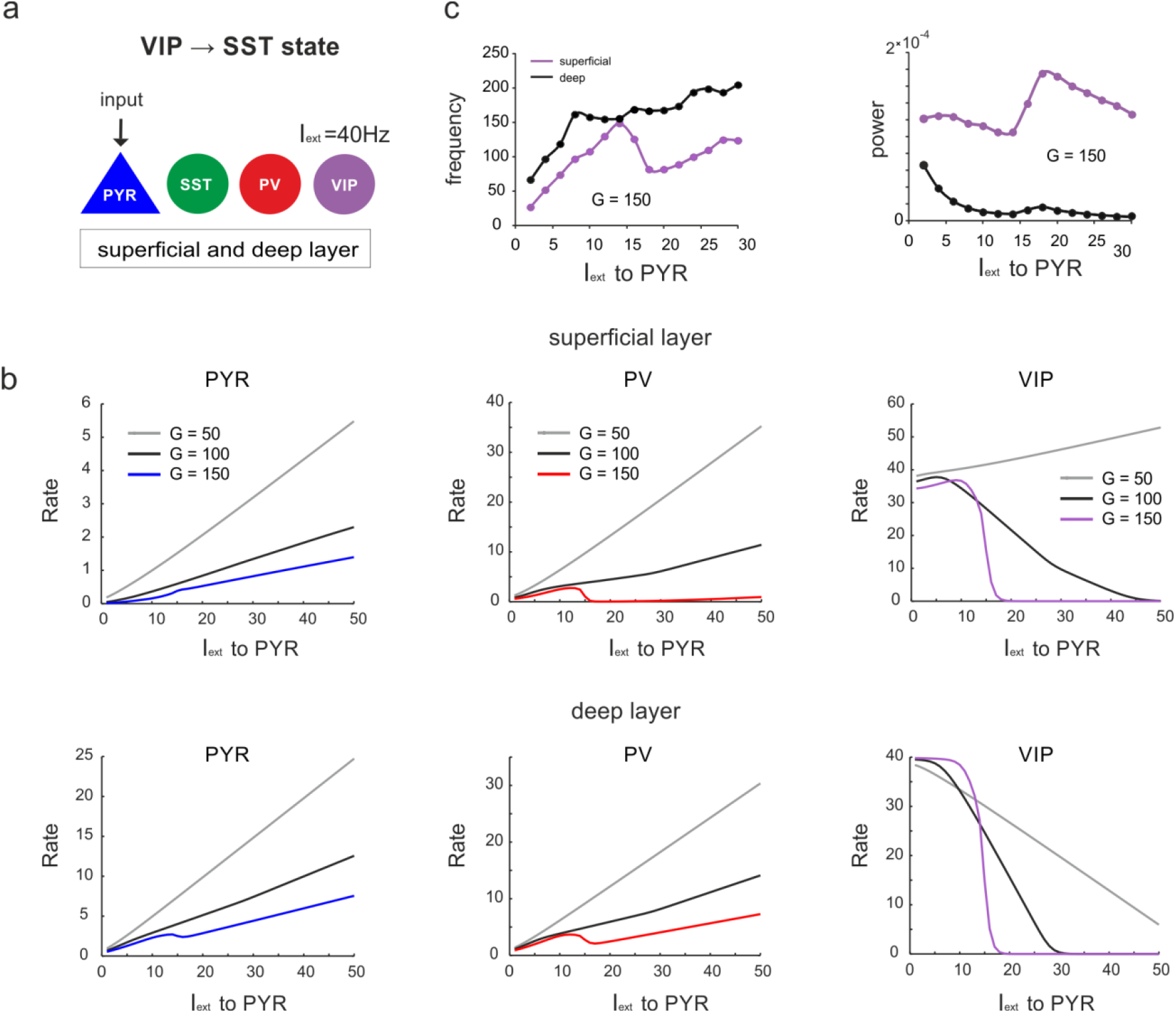
Response of different cell types to PYR cell input during the VIP dominated state. **a)** Superficial and deep PY cells were driven with a variable input, while a constant input of I_ext_ = 40 was given to VIP cells. **b)** Peak frequency and power in both layers as a function of PYR drive for an exemplary *G* value. **c)** Response of PYR, PV and VIP cells to PYR input for different values of *G* in superficial and deep layers.

**Supplementary Figure 9.**
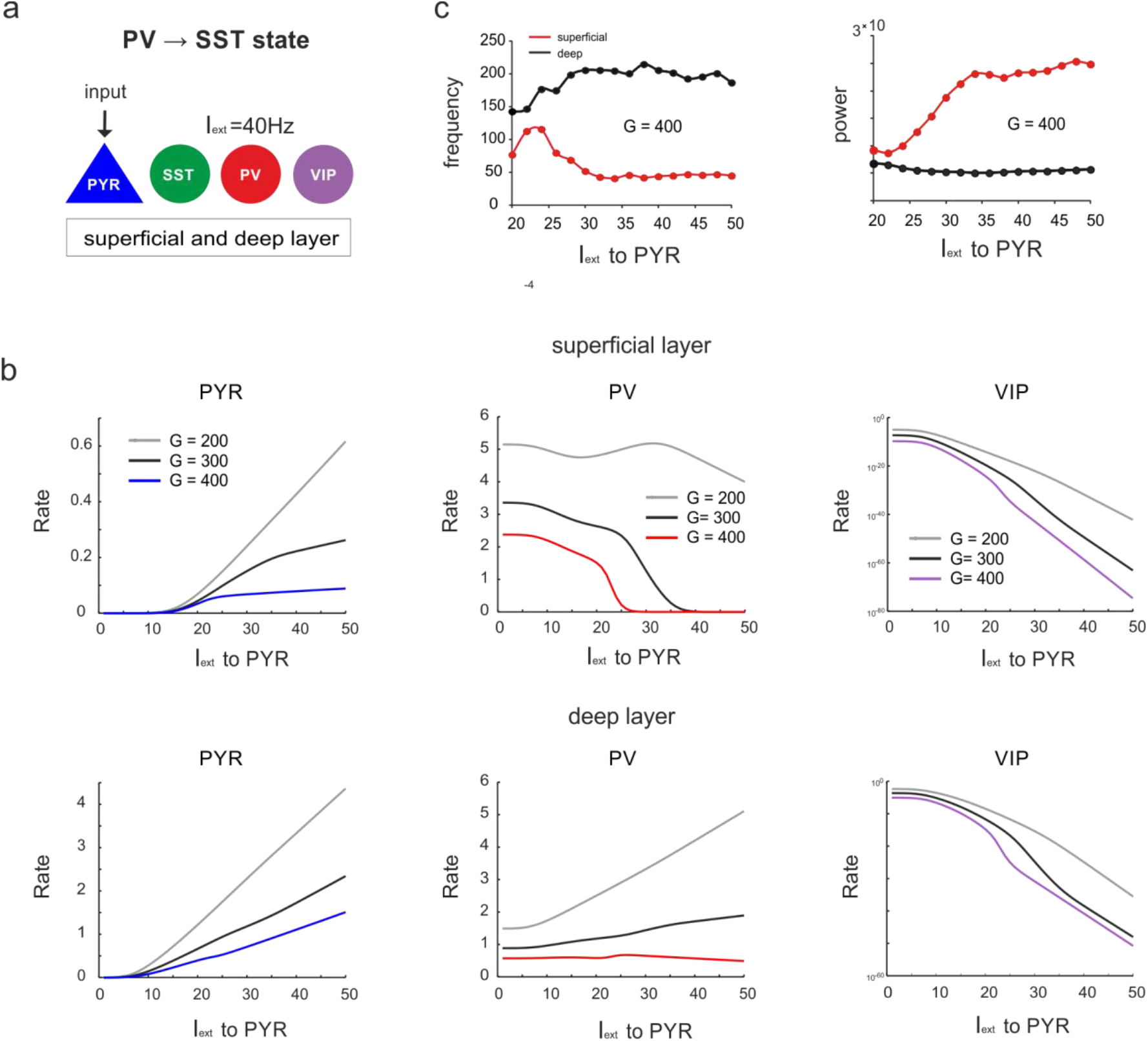
Response of different cell types to PYR cell input during the PV dominated state **a)** Superficial and deep PY cells were driven with a variable input, while a constant input of I_ext_ = 40 was given to PV cells. **b)** Peak frequency and power in both layers as a function of PYR drive for an exemplary *G* value. **c)** Response of PYR, PV and VIP cells to PYR input for different values of *G* in superficial and deep layers.

**Supplementary Figure 10.**
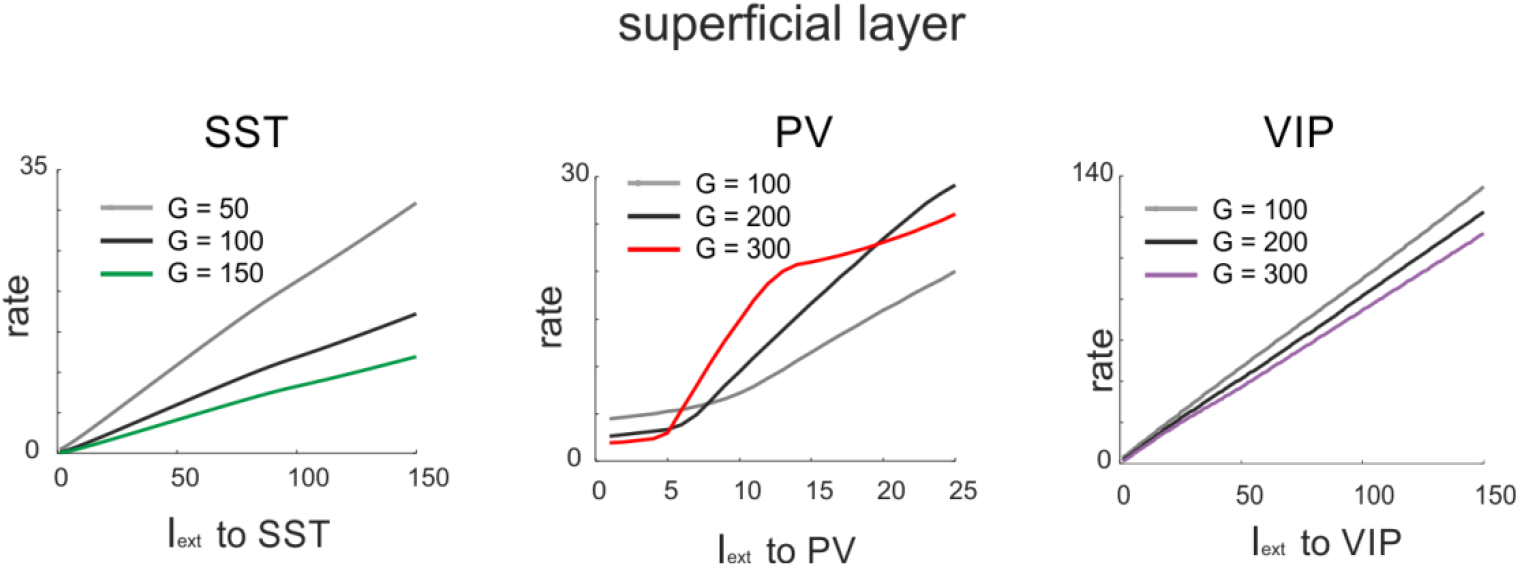
Response of SST, PV and VIP cells to input to the same cell type for three different values of *G* in the superficial layer.

**Supplementary Figure 11.**
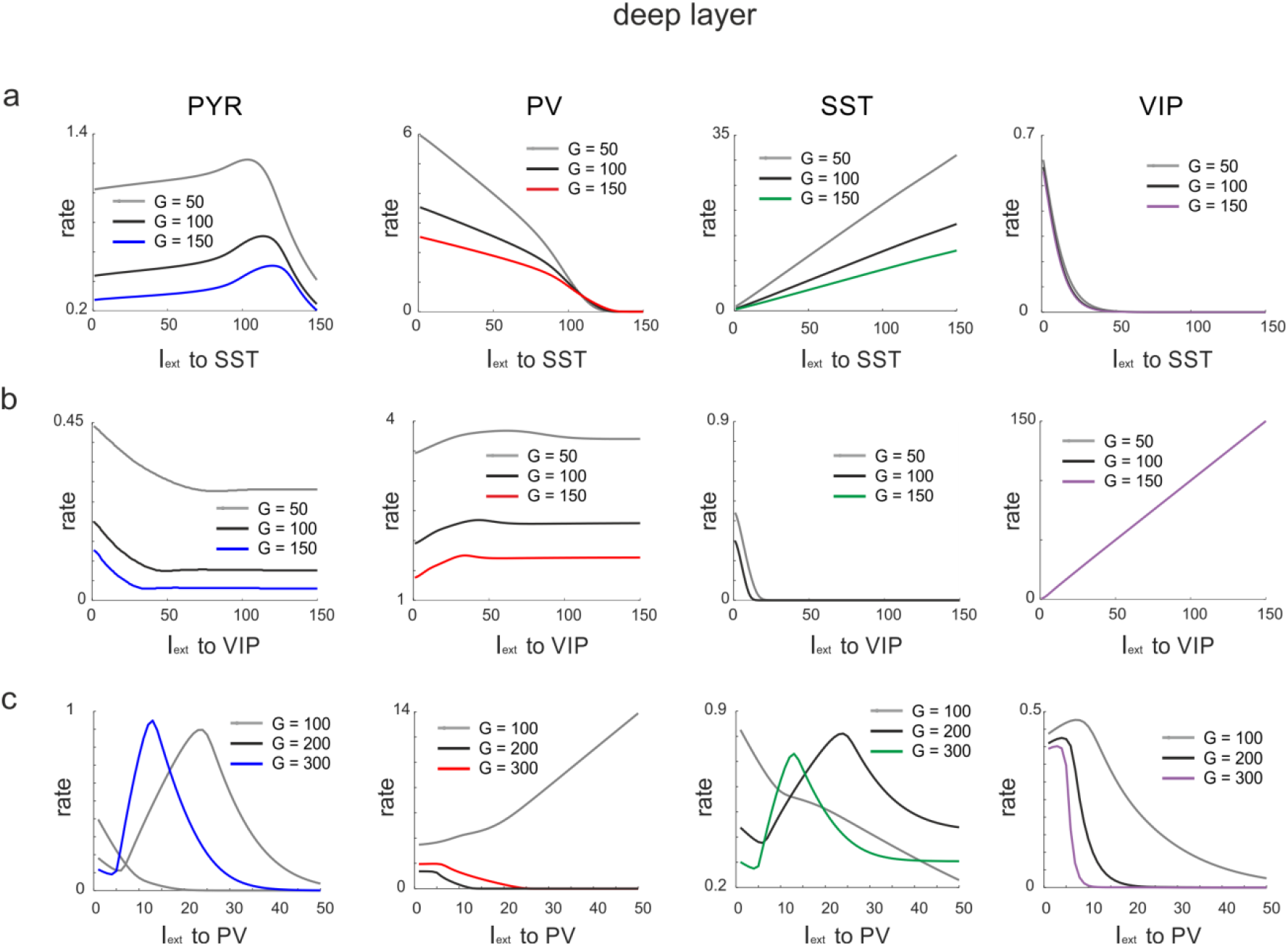
Responses to input to different cells after exchange between PV and SST recurrent connectivity. **a)** Response of all cell types to SST input in the deep layer for different values of *G*. Recurrent connectivity between PV cells was removed and enhanced between SST cells. **b)** Same as in a) for input to VIP cells. **c)** Same as in a) for input to PV cells.

**Supplementary Table 1.** Connection probability for different morphologically defined cell types, as described in Jiang et al. (2015). BC: basket cells; ChC: chandelier cells; MC: Martinotti cells; BTC: bitufted cells; SC: shrub cells; HEC: horizontally elongated cells; BPC: bipolar cells.

|  |  |  | presynaptic |  |  |  |  |  |  |  |  |  |  |
| --- | --- | --- | --- | --- | --- | --- | --- | --- | --- | --- | --- | --- | --- |
|  |  |  | L 2/3 |  |  |  |  | L 5 |  |  |  |  |  |
|  |  |  | PYR | BC | ChC | MC | BTC | PYR | BC | SC | HEC | MC | BPC |
| postsynaptic | L2/3 | PYR | 0.02 | 0.35 | 0.31 | 0.44 | 0.18 | --- | 0.1 | --- | --- | 0.21 | --- |
|  |  | BC | 0.19 | 0.47 | --- | 0.49 | 0.03 | 0.03 | 0.22 | --- | --- | 0.25 | --- |
|  |  | ChC | 0.19 | 0.18 | --- | --- | 0.26 | --- | 0.03 | --- | --- | --- | --- |
|  |  | MC | 0.2 | 0.11 | --- | 0.48 | 0.14 | --- | 0.03 | --- | --- | 0.3 | --- |
|  |  | BTC | --- | 0.14 | 0.38 | 0.46 | --- | --- | --- | --- | --- | 0.27 | --- |
|  | L5 | PYR | 0.04 | 0.06 | --- | 0.08 | --- | 0.02 | 0.25 | 0.1 | 0.3 | 0.21 | --- |
|  |  | BC | 0.08 | 0.14 | --- | 0.05 | --- | 0.11 | 0.48 | 0.03 | --- | 0.35 | --- |
|  |  | SC | 0.11 | 0 | --- | 0 | --- | 0.08 | 0.16 | --- | --- | --- | 0.31 |
|  |  | HEC | --- | 0.04 | --- | 0.44 | --- | --- | --- | --- | --- | 0.31 | --- |
|  |  | MC | --- | 0.02 | --- | --- | --- | --- | 0.13 | 0.63 | --- | 0.34 | --- |
|  |  | BPC | 0.08 | 0.04 | --- | --- | --- | 0.17 | 0.04 | --- | 0.5 | 0.33 | --- |

**Supplementary Table 2.** EPSP/IPSP strength for different morphologically defined cell types, as described in Jiang et al. (2015). BC: basket cells; ChC: chandelier cells; MC: Martinotti cells; BTC: bitufted cells; SC: shrub cells; HEC: horizontally elongated cells; BPC: bipolar cells.

|  |  |  | presynaptic |  |  |  |  |  |  |  |  |  |  |
| --- | --- | --- | --- | --- | --- | --- | --- | --- | --- | --- | --- | --- | --- |
|  |  |  | L 2/3 |  |  |  |  | L 5 |  |  |  |  |  |
|  |  |  | PYR | BC | ChC | MC | BTC | PYR | BC | SC | HEC | MC | BPC |
| postsynaptic | L2/3 | PYR | 0.3 | -0.5 | -0.35 | --- | -0.3 | --- | --- | -0.4 | --- | --- | -0.25 |
|  |  | BC | 1.6 | -0.7 | --- | --- | -0.2 | --- | 1 | -0.5 | --- | --- | -0.4 |
|  |  | ChC | 0.9 | -0.9 | --- | --- | -0.3 | --- | --- | -0.2 | --- | --- | 0 |
|  |  | MC | 1.3 | -0.4 | --- | --- | -0.4 | --- | --- | -0.3 | --- | --- | -0.7 |
|  |  | BTC | 0 | -0.69 | -0.47 | --- | --- | --- | --- | --- | --- | --- | -0.54 |
|  | L5 | PYR | 1.15 | -0.56 | --- | --- | --- | --- | 0.53 | -0.43 | --- | -0.81 | -0.45 |
|  |  | BC | 1.1 | -0.2 | --- | --- | --- | --- | 0.3 | -0.8 | -0.44 | -0.9 | -0.3 |
|  |  | SC | 1.3 | -0.8 | --- | --- | --- | --- | 1.2 | -1.2 | -0.53 | --- | -0.4 |
|  |  | HEC | 0.5 | --- | --- | -0.3 | --- | -0.3 | 0.5 | -0.4 | --- | --- | 0 |
|  |  | MC | --- | -0.4 | --- | --- | --- | --- | --- | --- | --- | --- | -0.4 |
|  |  | BPC | --- | -0.25 | --- | --- | --- | --- | --- | -0.6 | -1.4 | --- | -0.48 |

**Supplementary Table 3.** Morphological interneuron types, their genetic marker and proportion.

| Name | Genetic Marker | Proportion |
| --- | --- | --- |
| Basket Cell | PV | L2/3: 40% L5: 32% |
| Chandelier Cell | PV | L2/3: 2% |
| Shrub Cell | PV | L5: 18% |
| Horizontally-Elongated Cell | PV | L5: 10% |
| Martinotti Cell | SST | L2/3: 11% L5:32% |
| Bipolar Cell | VIP | L2/3: 17% |
| Bitufted Cell | VIP | L2/3: 10% |

**Supplementary Table 4.** Connectivity matrix corrected for cell proportions and scaled up by g = 100.

|  | presynaptic |  |  |  |  |  |  |  |  |  |
| --- | --- | --- | --- | --- | --- | --- | --- | --- | --- | --- |
| postsynaptic |  |  | L 2/3 |  |  |  | L5 |  |  |  |
|  |  |  | PYR | PV | SST | VIP | PYR | PV | SST | VIP |
|  | L2/3 | PYR | 0.6 | -1.44 | -0.29 | -0.11 | --- | -0.26 | -0.34 | --- |
|  |  | PV | 30.4 | -2.7 | -0.54 | -0.01 | 3 | -0.7 | -0.64 | --- |
|  |  | SST | 17.1 | -1.3 | --- | -0.16 | --- | -0.04 | --- | --- |
|  |  | VIP | 26 | -0.35 | -0.53 | -0.11 | --- | -0.06 | -1.34 | --- |
|  | L5 | PYR | 4.4 | -0.1 | -0.04 | --- | 0.6 | -1.98 | -0.4 | --- |
|  |  | PV | 10.4 | -0.9 | -0.02 | --- | 13.2 | -7.73 | -0.9 | --- |
|  |  | SST | 5.5 | --- | --- | --- | 4 | -0.41 | --- | -0.32 |
|  |  | VIP | --- | -0.13 | -0.58 | --- | --- | --- | -0.79 | --- |

**Supplementary Table 5.**

| Time Constants | Value |
| --- | --- |
| $\tau_{\text{PYR}}$ | 3 ms |
| $\tau_{\text{PV}}$ | 7 ms |
| $\tau_{\text{SST}}$ | 30 ms |
| $\tau_{\text{VIP}}$ | 10 ms |

## References

1. Gouwens, N. W. et al. Classification of electrophysiological and morphological neuron types in the mouse visual cortex. Nat. Neurosci. 22, 1182–1195 (2019).

2. Huang, Z. J. & Paul, A. The diversity of GABAergic neurons and neural communication elements. Nat. Rev. Neurosci. 20, 563–572 (2019).

3. Ascoli, G. a et al. Petilla terminology: nomenclature of features of GABAergic interneurons of the cerebral cortex. Nat. Rev. Neurosci. 9, 557–68 (2008).

4. Markram, H., et al. Interneurons of the neocortical inhibitory system. Nat. Rev. Neurosci. 5, 793–807 (2004).

5. Tremblay, R., Lee, S. & Rudy, B. GABAergic Interneurons in the Neocortex: From Cellular Properties to Circuits. Neuron 91, 260–292 (2016).

6. Pfeffer, C. K., Xue, M., He, M., Huang, Z. J. & Scanziani, M. Inhibition of inhibition in visual cortex: The logic of connections between molecularly distinct interneurons. Nat. Neurosci. 16, 1068–1076 (2013).

7. Jiang, X. et al. Principles of connectivity among morphologically defined cell types in adult neocortex. Science (80-.). 350, (2015).

8. Kätzel, D., Zemelman, B. V, Buetfering, C., Wölfel, M. & Miesenböck, G. The columnar and laminar organization of inhibitory connections to neocortical excitatory cells. Nat. Neurosci. 14, 100–107 (2011).

9. Harris, K. D. & Shepherd, G. M. G. The neocortical circuit: themes and variations. Nat. Neurosci. 18, 170–181 (2015).

10. Cardin, J. A. Inhibitory Interneurons Regulate Temporal Precision and Correlations in Cortical Circuits. Trends Neurosci. 41, 689–700 (2018).

11. Douglas, R. J. & Martin, K. a. C. Neuronal Circuits of the Neocortex. Annu. Rev. Neurosci. 27, 419–451 (2004).

12. Beul, S. F. & Hilgetag, C. C. Towards a canonical agranular cortical microcircuit. Front. Neuroanat. 8, 1–8 (2015).

13. Miller, K. D. Canonical computations of cerebral cortex. (2016). doi:10.1016/j.conb.2016.01.008

14. Kepecs, A. & Fishell, G. Interneuron cell types are fit to function. Nature 505, 318–326 (2014).

15. Fishell, G. & Kepecs, A. Annual Review of Neuroscience Interneuron Types as Attractors and Controllers. Rev. Adv. first posted (2019). doi:10.1146/annurev-neuro-070918

16. Maffei, A. Fifty shades of inhibition. Current Opinion in Neurobiology 43, 43–47 (2017).

17. Letzkus, J. J., Wolff, S. B. E. & Lüthi, A. Disinhibition, a Circuit Mechanism for Associative Learning and Memory. Neuron 88, 264–276 (2015).

18. Pi, H. J. et al. Cortical interneurons that specialize in disinhibitory control. Nature 503, 521–524 (2013).

19. Walker, F. et al. Parvalbumin-and vasoactive intestinal polypeptide-expressing neocortical interneurons impose differential inhibition on Martinotti cells. Nat. Commun. 7, 13664 (2016).

20. Jackson, J., Ayzenshtat, I., Karnani, M. M. & Yuste, R. VIP+ interneurons control neocortical activity across brain states. J. Neurophysiol. 115, 3008–3017 (2016).

21. Fu, Y. et al. A cortical circuit for gain control by behavioral state. Cell 156, 1139–1152 (2014).

22. Cardin, J. et al. Driving fast-spiking cells induces gamma rhythm and controls sensory responses. Nature 459, 663–7 (2009).

23. Sohal, V., Zhang, F., Yizhar, O. & Deisseroth, K. Parvalbumin neurons and gamma rhythms enhance cortical circuit performance. Nature 459, 698–702 (2009).

24. Chen, G. et al. Distinct Inhibitory Circuits Orchestrate Cortical beta and gamma Band Oscillations Article Distinct Inhibitory Circuits Orchestrate Cortical beta and gamma Band Oscillations. Neuron 96, 1403–1418.e6 (2017).

25. Lee, J. H., Koch, C. & Mihalas, S. A Computational Analysis of the Function of Three Inhibitory Cell Types in Contextual Visual Processing. Front. Comput. Neurosci. 11, 1–15 (2017).

26. Lee, J. H. & Mihalas, S. Visual processing mode switching regulated by VIP cells. Sci. Rep. 7, 1–15 (2017).

27. Hertäg, L. & Sprekeler, H. Amplifying the redistribution of somato-dendritic inhibition by the interplay of three interneuron types. PLoS Comput. Biol. 15, 1–40 (2019).

28. Carlos, L., Yang, G. R., Mejias, J. F. & Wang, X. Paradoxical response reversal of top-down modulation in cortical circuits with three interneuron types. Elife 1–15 (2017).

29. Lee, B. et al. Combined Positive and Negative Feedback Allows Modulation of Neuronal Oscillation Frequency during Article Combined Positive and Negative Feedback Allows Modulation of Neuronal Oscillation Frequency during Sensory Processing. CellReports 25, 1548–1560.e3 (2018).

30. Bastos, A. M., Loonis, R., Kornblith, S., Lundqvist, M. & Miller, E. K. Laminar recordings in frontal cortex suggest distinct layers for maintenance and control of working memory. Proc. Natl. Acad. Sci. U. S. A. 201710323 (2018). doi:10.1073/pnas.1710323115

31. Adesnik, H. Layer-specific excitation/inhibition balances during neuronal synchronization in the visual cortex. J. Physiol. 596, 1639–1657 (2018).

32. van Kerkoerle, T. et al. Alpha and gamma oscillations characterize feedback and feedforward processing in monkey visual cortex. Proc. Natl. Acad. Sci. 111, 14332–14341 (2014).

33. Veit, J., Hakim, R., Jadi, M. P., Sejnowski, T. J. & Adesnik, H. Cortical gamma band synchronization through somatostatin interneurons. Nat. Neurosci. 20, 951–959 (2017).

34. Jiang, X. et al. Principles of connectivity among morphologically defined cell types in adult neocortex. Science (80-.). 350, 1–51 (2015).

35. Deco, G. et al. Resting-state functional connectivity emerges from structurally and dynamically shaped slow linear fluctuations. J. Neurosci. 33, 11239–52 (2013).

36. Jobst, B. M. et al. Increased Stability and Breakdown of Brain Effective Connectivity During Slow-Wave Sleep: Mechanistic Insights from Whole-Brain Computational Modelling. Sci. Rep. 7, 4634 (2017).

37. Sakata, S. & Harris, K. Laminar structure of spontaneous and sensory-evoked population activity in auditory cortex. Neuron 64, 404–18 (2009).

38. Senzai, Y., Fernandez-Ruiz, A. & Buzsáki, G. Layer-Specific Physiological Features and Interlaminar Interactions in the Primary Visual Cortex of the Mouse. Neuron 101, 500–513.e5 (2019).

39. Haegens, X. S. et al. Laminar Profile and Physiology of the Rhythm in Primary Visual, Auditory, and Somatosensory Regions of Neocortex. J. Neurosci. 35, 14341–14352 (2015).

40. Buffalo, E. A., Fries, P., Landman, R., Buschman, T. J. & Desimone, R. Laminar differences in gamma and alpha coherence in the ventral stream. Proc. Natl. Acad. Sci. 108, 11262–11267 (2011).

41. Bonaiuto, J. J. et al. Lamina-specific cortical dynamics in human visual and sensorimotor cortices. Elife 7, (2018).

42. Pluta, S. et al. A direct translaminar inhibitory circuit tunes cortical output. Nat. Neurosci. 18, 1–11 (2015).

43. Ferrell JE. Ultrasensitivity I. 39, 496–503 (2015).

44. Jr, J. E. F. Self-perpetuating states in signal transduction: positive feedback, double-negative feedback and bistability. 140–148 doi:10.1016/S0955-0674(02)00314-9

45. Adesnik, H., Bruns, W., Taniguchi, H. & Huang, Z. J. A Neural Circuit for Spatial Summation in Visual Cortex. Nature 490, 226–231 (2012).

46. Dipoppa, M. et al. Vision and Locomotion Shape the Interactions between Neuron Types in Mouse Visual Cortex. Neuron 98, 602–615.e8 (2018).

47. Gieselmann, M. A. & Thiele, A. Comparison of spatial integration and surround suppression characteristics in spiking activity and the local field potential in macaque V1. Eur. J. Neurosci. 28, 447–459 (2008).

48. Hakim, R., Shamardani, K. & Adesnik, H. A neural circuit for gamma-band coherence across the retinotopic map in mouse visual cortex. Elife 7, 1–17 (2018).

49. Fino, E. & Yuste, R. Dense Inhibitory Connectivity in Neocortex. Neuron 69, 1188–1203 (2011).

50. Urban-Ciecko, J. & Barth, A. L. Somatostatin-expressing neurons in cortical networks. Nat. Rev. Neurosci. 17, 401–409 (2016).

51. Naka, A. et al. Complementary networks of cortical somatostatin interneurons enforce layer specific control. Elife 8, (2019).

52. Ferguson, B. R. & Gao, W. J. Pv interneurons: critical regulators of E/I balance for prefrontal cortex-dependent behavior and psychiatric disorders. Frontiers in Neural Circuits 12, 37 (2018).

53. Garcia-Junco-Clemente, P. et al. An inhibitory pull-push circuit in frontal cortex. Nat. Neurosci. 20, 389–392 (2017).

54. Wang, X. J. & Yang, G. R. A disinhibitory circuit motif and flexible information routing in the brain. Curr. Opin. Neurobiol. 49, 75–83 (2018).

55. Bartos, M., Vida, I. & Jonas, P. Synaptic mechanisms of synchronized gamma oscillations in inhibitory interneuron networks. Nat. Rev. Neurosci. 8, 45–56 (2007).

56. Hahn, G., Bujan, A. F., Frégnac, Y., Aertsen, A. & Kumar, A. Communication through Resonance in Spiking Neuronal Networks. PLoS Comput. Biol. 10, e1003811 (2014).

57. Tyson, J. J., Chen, K. C. & Novak, B. Sniffers, buzzers, toggles and blinkers: dynamics of regulatory and signaling pathways in the cell. 221–231 (2003). doi:10.1016/S0955-0674(03)00017-6

58. Markov, N. T. et al. Anatomy of hierarchy: Feedforward and feedback pathways in macaque visual cortex. J. Comp. Neurol. 522, 225–259 (2014).

59. Markov, N. T. et al. Cortical high-density counterstream architectures. Science (80-.). 342, 1238406 (2013).

60. Christophel, T. B., Klink, P. C., Spitzer, B., Roelfsema, P. R. & Haynes, J. D. The Distributed Nature of Working Memory. Trends Cogn. Sci. 21, 111–124 (2017).

61. Ardid, S., Wang, X.-J. & Compte, A. An integrated microcircuit model of attentional processing in the neocortex. J. Neurosci. 27, 8486–8495 (2007).

62. Dehaene, S., Sergent, C. & Changeux, J.-P. A neuronal network model linking subjective reports and objective physiological data during conscious perception. Proc. Natl. Acad. Sci. U. S. A. 100, 8520–5 (2003).

63. Dehaene, S. & Changeux, J.-P. Experimental and theoretical approaches to conscious processing. Neuron 70, 200–27 (2011).

64. van Vugt, B. et al. The threshold for conscious report: Signal loss and response bias in visual and frontal cortex. Science (80-.). 360, 537–542 (2018).

65. Kunze, T., Peterson, A. D. H., Haueisen, J. & Knösche, T. R. A model of individualized canonical microcircuits supporting cognitive operations. PLoS One 12, e0188003 (2017).

66. Nelson, M. J. et al. Neurophysiological dynamics of phrase-structure building during sentence processing. Proc. Natl. Acad. Sci. U. S. A. 114, E3669–E3678 (2017).

67. Palliera, C., Devauchellea, A. D. & Dehaenea, S. Cortical representation of the constituent structure of sentences. Proc. Natl. Acad. Sci. U. S. A. 108, 2522–2527 (2011).

68. Compte, A., Brunel, N., Goldman-Rakic, P. S. & Wang, X.-J. Synaptic Mechanisms and Network Dynamics Underlying Spatial Working Memory in a Cortical Network Model. Cereb. Cortex 10, 910–923 (2000).

69. Kim, Y. et al. Brain-wide Maps Reveal Stereotyped Cell-Type-Based Cortical Architecture and Subcortical Sexual Dimorphism. Cell 171, 456–469.e22 (2017).

70. Fulcher, B. D., Murray, J. D., Zerbi, V. & Wang, X.-J. Multimodal gradients across mouse cortex. Proc. Natl. Acad. Sci. U. S. A. 116, 4689–4695 (2019).

71. Demirtaş, M. et al. Hierarchical Heterogeneity across Human Cortex Shapes Large-Scale Neural Dynamics. Neuron 101, 1181–1194.e13 (2019).

72. Silberberg, G. & Markram, H. Disynaptic Inhibition between Neocortical Pyramidal Cells Mediated by Martinotti Cells. Neuron 53, 735–746 (2007).

73. Brunel, N. & Wang, X. What determines the frequency of fast network oscillations with irregular neural discharges? I. Synaptic dynamics and excitation-inhibition balance. J. Neurophysiol. 90, 415–30 (2003).

74. Jia, X., Xing, D. & Kohn, A. No consistent relationship between gamma power and peak frequency in macaque primary visual cortex. J. Neurosci. 33, 17–25 (2013).

75. Zhang. Long-range and local circuits for top-down modulation of visual cortex processing. Science (80-.). 6263–6263 (2014).

76. Gonchar, Y. & Burkhalter, A. Distinct GABAergic Targets of Feedforward and Feedback Connections between Lower and Higher Areas of Rat Visual Cortex. J. Neurosci. 23, 10904–10912 (2003).

77. Ray, S. & Maunsell, J. H. R. Differences in gamma frequencies across visual cortex restrict their possible use in computation. Neuron 67, 885–96 (2010).

78. Pakan, J. M. et al. Behavioral-state modulation of inhibition is context-dependent and cell type specific in mouse visual cortex. Elife 5, (2016).

79. Hahn, G., Ponce-Alvarez, A., Deco, G., Aertsen, A. & Kumar, A. Portraits of communication in neuronal networks. Nat. Rev. Neurosci. (2018). doi:10.1038/s41583-018-0094-0

80. Klimesch, W., Sauseng, P. & Hanslmayr, S. EEG alpha oscillations: The inhibition-timing hypothesis. Brain Res. Rev. 53, 63–88 (2007).

81. Jensen, O. & Mazaheri, A. Shaping functional architecture by oscillatory alpha activity: gating by inhibition. Front. Hum. Neurosci. 4, 186 (2010).

82. Singer, W. Neuronal synchrony: a versatile code for the definition of relations? Neuron 24, 49–65, 111–25 (1999).

83. Fries, P. Rhythms for Cognition: Communication through Coherence. Neuron 88, 220–235 (2015).

84. Wilson, H. & Cowan, J. Excitatory and inhibitory interactions in localized populations of model neurons. Biophys. J. 12, 1–24 (1972).

85. Hyafil, A., Fontolan, L., Kabdebon, C., Gutkin, B. & Giraud, A. L. Speech encoding by coupled cortical theta and gamma oscillations. Elife 4, 1–45 (2015).

86. Vierling-Claassen, D., Cardin, J. A., Moore, C. I. & Jones, S. R. Computational modeling of distinct neocortical oscillations driven by cell-type selective optogenetic drive: separable resonant circuits controlled by low-threshold spiking and fast-spiking interneurons. Front. Hum. Neurosci. 4, 198 (2010).

87. Mejias, J. F., Murray, J. D., Kennedy, H. & Wang, X.-J. Feedforward and feedback frequency-dependent interactions in a large-scale laminar network of the primate cortex. Science (80-.). 065854 (2016). doi:10.1101/065854

88. Jr, J. E. F. & Machleder, E. M. The biochemical basis of an all or none switch in xenopus oocytes. 280, 895–898 (1998).

